# Computational and biochemical analyses reveal that cofilin-2 self assembles into amyloid-like structures and promotes the aggregation of other proteinaceous species: Pathogenic relevance to myopathies

**DOI:** 10.1101/2021.11.27.470221

**Authors:** Vibha Kaushik, Eva-Maria Hanschmann, Daniela Brünnert, Kumari Prerna, Bibin G. Anand, Phulwanti Kumari Sharma, Karunakar Kar, Pankaj Goyal

## Abstract

Cofilin-2 is a member of the ADF/cofilin family, expressed extensively in adult muscle cells and involved in muscle maintenance and regeneration. Phosphorylated cofilin-2 is found in pre-fibrillar aggregates formed during idiopathic dilated cardiomyopathy. A recent study shows that phosphorylated cofilin-2, under oxidative distress, forms fibrillar aggregates. However, it remains unknown if cofilin-2 holds an innate propensity to form amyloid-like structures. In the present study, we employed various computational and biochemical techniques to explore the amyloid-forming potential of cofilin-2. We report that cofilin-2 possesses aggregation-prone regions (APRs), and these APRs get exposed to the surface, become solvent-accessible, and are involved in the intermolecular interactions during dimerization, an early stage of aggregation. Furthermore, the cofilin-2 amyloids, formed under physiological conditions, are capable of cross-seeding other monomeric globular proteins and amino acids, thus promoting their aggregation. We further show that Cys-39 and Cys-80 are critical in maintaining the thermodynamic stability of cofilin-2. The destabilizing effect of oxidation at Cys-39 but not that at Cys-80 is mitigated by Ser-3 phosphorylation. Cysteine oxidation leads to partial unfolding and loss of structure, suggesting that cysteine oxidation further induces early events of cofilin-2 aggregation. Overall, our results pose a possibility that cofilin-2 amyloidogenesis might be involved in the pathophysiology of diseases, such as myopathies. We propose that the exposure of APRs to the surface could provide mechanistic insight into the higher-order aggregation and amyloidogenesis of cofilin-2. Moreover, the cross-seeding activity of cofilin-2 amyloids hints towards its involvement in the hetero-aggregation in various amyloid-linked diseases.

## 1. Introduction

Cofilin-2 belongs to the ADF/cofilin family of actin-dynamizing proteins, including two other members, i.e., ADF, expressed in endothelial and epithelial cells, and cofilin-1 expressed ubiquitously in all adult tissues and embryonic muscle cells [1]. Despite highly identical sequences (∼80% sequence identity) and similar affinities for G-actin and F-actin between the cofilins, cofilin-2 severs actin filaments more efficiently than cofilin-1 [2,3]. Moreover, cofilin-2 possesses only two cysteine residues, i.e., Cys-39 and Cys-80, while cofilin-1 has two additional cysteines at positions 139 and 147 [4]. Cofilin-2 has evolved its biochemical and cellular properties to regulate actin dynamics in sarcomeres [5]. It plays an essential role in mediating cell cycle progression, proliferation, and myogenic differentiation in C2C12 myoblasts [6] and is critically involved in muscle maintenance [7]. The activity of cofilin-2 is under tight regulation of phosphorylation, dephosphorylation, presence of calcium, environmental pH, and interaction with muscle Lim protein [8,9], phosphatidylinositol 4,5-bisphosphate (PIP_2_) [10], and pro-proliferative miRNAs [11].

It is now well known that oxidation of the cysteine residues in cofilin-1 under pathophysiological conditions, such as oxidative distress, is detrimental for its actin-binding and severing activity, thus impairing cell migration [12]. Moreover, the cofilin-saturated actin forms actin-cofilin rods under oxidative distress, wherein Cys-39 and Cys-147 of cofilin-1 form intermolecular disulfide bonds [13]. These rods are reported to worsen the pathology of amyloid-linked neurodegenerative diseases, such as Alzheimer’s disease [14]. Myofibrillar accumulation of amyloid-like fibrils and their precursor seeds have been reported in the hearts of idiopathic dilated cardiomyopathy (iDCM) patients [15,16]. Furthermore, Subramanian *et al*. showed that hyperphosphorylated cofilin-2, along with its interactome, actin, and MLC-II, is a component of the pre-amyloid oligomer aggregates in the myocardium of these patients, suggesting that the perturbed cofilin-2 promotes the pathogenesis of iDCM [17].

Apart from the involvement of cofilins in protein aggregation and related diseases, studies revealed that cofilin-1 forms oligomeric species *in vitro* [18]. Our group showed that cofilin-1 holds an innate tendency of oligomerization and exists both as oligomers and monomers in platelets and endothelial cells under physiological conditions. Cofilin-1 oligomerization increases during platelet activation with thrombin and is negatively regulated via phosphorylation [19]. Our recent finding revealed that cofilin-1 possesses an intrinsic propensity to form higher-order amyloidogenic fibrillar structures, and this amyloidogenesis is further aggravated by Cys-80 oxidation [20]. It has been divulged that the phosphorylated form of cofilin-2, under oxidative distress, forms fibrillar aggregates, which involves Cys-39 [4]. However, it is not yet clear whether cofilin-2 has an intrinsic property of amyloidogenesis under physiological conditions.

Therefore, we explored the innate propensity of cofilin-2 to form amyloid-like structures under physiological conditions and the effect of oxidation and phosphorylation on its amyloidogenesis. The current study suggests that cofilin-2 has an inherent tendency of amyloid formation, and it does not necessarily require any post-translational modification to form these higher-order species. We also suggest that cofilin-2 amyloids can promote the aggregation of globular proteins and amino acids. Our results provide a possible link between cofilin-2 amyloid formation and the pathology of diseases, such as myopathies.

### 2. Materials and methods

### 2.1. Identification of the intrinsic amyloidogenic potential of cofilin-2

Cofilin-2 was analyzed for its inherent amyloidogenic property, both at the primary sequence and tertiary structural levels. For the primary sequence analysis, the protein sequence retrieved from the NCBI protein database with accession number NP_6195791 was submitted in the required format to the machine learning-based open-access predictors viz., AGGRESCAN [21], TANGO [22], FoldAmyloid [23], and AmylPred 2 [24].

For tertiary structural analysis, the cofilin-2 was modeled and validated (Supplementary information SI-1, Figure S1). The modeled structure was then submitted to a structure-based prediction tool, Aggrescan3D 2.0, that considers the conformational aspects and surface exposure of residues in a folded protein to identify [25] the aggregation-prone residues. The analysis was performed at a sphere radius of 5 Å in dynamic mode.

### 2.2. Cofilin-2 homodimerization, molecular dynamics simulations, and solvent accessibility analysis

The homodimerization of cofilin-2 was performed employing Z-Dock-3.0.2f+IRaPPA re-ranking version [26,27] by providing the 3D structure of cofilin-2, both as ligand and receptor. The top-10 dimer complexes were subjected to molecular dynamics study by Gromacs 2019 using the OPLS-AA/L force field in a cubic box with SPC water molecules as the solvent and 0.1 M NaCl to neutralize the system. The energy of the system was minimized by the Steepest Descent method. Further, the solvent was equilibrated around the protein in two phases, NVT and NPT, for 100 ps each to stabilize the temperature and pressure, respectively. Later, the density was also stabilized. After the system got stabilized, molecular dynamics was run for 100 ns. Later, the solvent accessibility of APRs in homodimers was calculated and the critical residues involved in the intermolecular interactions were identified using the BIOVIA Discovery Studio 2020 Client visualizer (Dassault Systèmes^®^, France).

### 2.3. Cloning of human *cofilin-2* gene, expression, and purification of the protein

As described previously [20], the human cofilin-2 gene was cloned in the pET 15b plasmid using specific oligonucleotides (Forward: **CATATG**gcttctggagttacag; Reverse: **GGATCC**ttataagggttttccttcaagtg) between the Nde1 and BamH1 restriction enzyme sites with an N-terminal 6X His-tag. The recombinant plasmid was then transformed to the Rosetta2(DE3)pLysS *E*.*coli* cells. These cells were then grown further at 37 °C until OD_600_ reached 0.6-0.8. Later, the promoter of the recombinant plasmid was induced with 0.3 mM IPTG, and the cells were further grown at 16 °C for 24 h to express human cofilin-2. The induced cells were centrifuged at 4500 g; the pellet was weighed and resuspended in ice-cold lysis buffer (1X PBS with 200 μM PMSF and 50 μg/ml lysozyme; pH 7.4), as per its weight, and incubated on ice for 30 min. Cell lysis was performed by sonication, the lysate was centrifuged at 24,000 g for 40 min at 4 °C, and the soluble fraction was filtered. Further, a column was packed using HisPur^™^ Ni-NTA resins, equilibrated with 1X PBS, and the filtered soluble fraction was then loaded onto the column. The column was then washed thoroughly with 3 column volumes of 1X PBS. The His-tagged cofilin-2 was eluted with elution buffer (1X PBS containing 250 mM imidazole and 10% glycerol; pH 7.4). The purity of the eluted cofilin-2 was analyzed by SDS-PAGE and western blotting, while Bradford’s assay was employed to assess the protein concentration. A protein yield of ∼9 mg/l of culture with > 95% purity was obtained. The purified protein was then desalted against 1X PBS, pH 7.4, using Bio-Rad-Econo-Pac 10DG desalting columns.

### 2.4. Native (non-denaturing) polyacrylamide gel electrophoresis (PAGE)

The aggregation reaction was initiated by incubating 25 μM and 50 μM of the purified cofilin-2 at 37 °C, shaking at 1200 rpm for 24 h on a thermomixer. Further, the incubated cofilin-2 samples were assessed by Native-PAGE to confirm the formation of higher-order entities after the aggregation process, as described previously [20]. Briefly, the samples were loaded onto a 12% Tris-HCl polyacrylamide gel and electrophoresed using the mini-PROTEIN II electrophoresis system (Bio-Rad, USA) at 25 Amp and 4 °C throughout the process. The gel was developed by silver staining and visualized using a gel documentation unit (Bio-Rad USA).

### 2.5. Congo Red spectral shift assay

The aggregated cofilin-2 samples at concentrations of 25 μM and 50 μM were incubated with a final concentration of 35 μM Congo Red (CR) for 5 min at room temperature. The absorption spectra were then recorded between 400 nm to 600 nm using a plate spectrophotometer (Thermo Scientific, Multiskan GO, USA).

### 2.6. Thioflavin T binding assay and fluorescence microscopy

The cofilin-2 samples were mixed with a final concentration of 20 μM Thioflavin T (ThT) and incubated in the dark for 5 min at room temperature. The fluorescence spectra were then recorded using the LS-55 fluorescence spectrophotometer (Perkin Elmer, USA) from 470 nm to 700 nm after exciting the dye at 450 nm. The excitation and emission slit widths were adjusted to 10 nm and 5 nm, respectively, with a path length of 1 cm.

The morphology of the cofilin-2 aggregates was analyzed by fluorescence microscopy. The aggregated sample was drop cast on a glass slide, air-dried, stained with 1.25 mM of ThT, and was then visualized using the ECLIPSE 90i fluorescence microscope (Nikon, Japan) at 20X magnification.

### 2.7. Cross-seeding of globular proteins and amino acids by cofilin-2 amyloid seeds

For analyzing the cross-seeding activity of cofilin-2, the cofilin-2 amyloids were centrifuged at maximum speed (386,000 g, 30 min), and the fibrillar entities were isolated as a pellet. We then added 5% w/w cofilin-2 seeds to a mixture of globular protein monomers containing BSA, lysozyme, insulin, myoglobin, and cytochrome c, at an equimolar concentration of 2 μM for each protein. Similarly, for analyzing the cross-seeding effect on the amino acids, a mixture of amino acids at an equimolar concentration of 33 μM each of Ala, Glu, Arg, Trp, Tyr, and Phe was spiked with cofilin-2 amyloid seeds. Both the sets were incubated at 37 °C. Later, the aggregation was monitored by native PAGE, ThT binding assay, and fluorescence microscopy, as mentioned above.

### 2.8. Stability assessment of cofilin-2 oxidation and phosphorylation mimics

The 3D structure of cofilin-2 was submitted to the Aggrescan3D 2.0 portal, and the two cysteines, Cys-39 and Cys-80, were replaced with Asp, separately for single mutations, and simultaneously to create a double oxidation mimic. For mimicking phosphorylation, Ser-3 was replaced with Asp. The mutants were then analyzed at a distance of aggregation known as the sphere radius of 5 Å employing the CABS-flex approach [28] that enables fast simulation of near-native dynamics of globular protein. The stability of the cofilin-2 mutants with reference to the wild type (WT) protein was assessed in terms of the energy differences calculated by the FoldX algorithm [29].

### 2.9. *In silico* mutagenesis, MD simulations, and secondary structure analysis

The cofilin-2 modeled structure was subjected to *in silico* mutagenesis at its cysteine residues using Schr□dinger suite 2018-1 (Schr□dinger, USA). The cofilin-2 WT structure was prepared by the protein preparation wizard of Maestro with default settings. Later, the cysteine residues were replaced with aspartic acid (C to D) mutations for mimicking oxidation. Additionally, the serine residue at position 3 was mutated to aspartic acid (S3D) to generate the phospho-mimic under the residue and loop mutation panel of the Bioluminate interface. Further, we minimized the energy of the mutant structures in an implicit solvent environment and placed them in a 10 Å x 10 Å x 10 Å-dimensioned orthorhombic box, solvated with SPC water molecules. The system was then neutralized with 0.15 M NaCl and sodium or chloride ions, as required. The system was then subjected to the OPLS3e force field, and a 10 ns molecular dynamics simulation was performed by employing the Desmond molecular dynamics module [30] at the temperature and pressure conditions of 300 K and 1.01 bar, respectively. Later, the Secondary Structure Elements (SSE) of the wild type and mutant structures were analyzed under the simulation interaction diagram panel. The BIOVIA Discovery Studio 2020 Client visualizer (Dassault Systèmes^®^, France) was used for visualizing the structures.

## 3. Results

### 3.1. Cofilin-2 holds an innate property of amyloidogenesis

We used four well-established web-based amyloidogenic prediction tools (AGGRESCAN, TANGO, FoldAmyloid, and AmylPred 2) to analyze the primary sequence of cofilin-2 for its inherent tendency to aggregate. The short stretches with high aggregation propensity predicted in a protein sequence by these tools are termed Aggregation Prone Regions (APRs). AGGRESCAN predicted nine APRs in cofilin-2; A_35_-L_40_, Q_46_-E_50_, Q_54_-D_59_, T_69_-L_75_, Y_82_-A_87_, L_99_-A_105_, S_113_-A_118_, K_126_-G_130_, and G_155_-L_161_. TANGO identified four APRs (A_35_-S_41_, Y_68_-V_72_, L_99_-W_104_, and N_156_-L_161_). FoldAmyloid predicted five APRs (V_36_-L_40_, Q_54_-G_58_, V_72_-P_76_, R_81_-A_87_, and L_99_-A_105_). AmylPred 2 determined seven APRs in cofilin-2 (I_12_-F_15_, A_35_-L_40_, I_47_-V_49_, Q_54_-D_59_, S_70_-L_75_, D_98_-A_105_, and G_155_-S_160_) (Figure 1A and Table S1). Cumulatively, ten regions in the cofilin-2 sequence were predicted. Two regions were commonly identified by at least three tools (Figure 1A, dotted boxes) and two stretches by at least two tools (Figure 1A, dashed boxes). Additionally, AGGRESCAN predicted two unique regions, and AmylPred 2 uniquely identified one short region. All the tools predicted three regions in consensus for cofilin-2 (Figure 1A, solid boxes).

**Figure 1:**
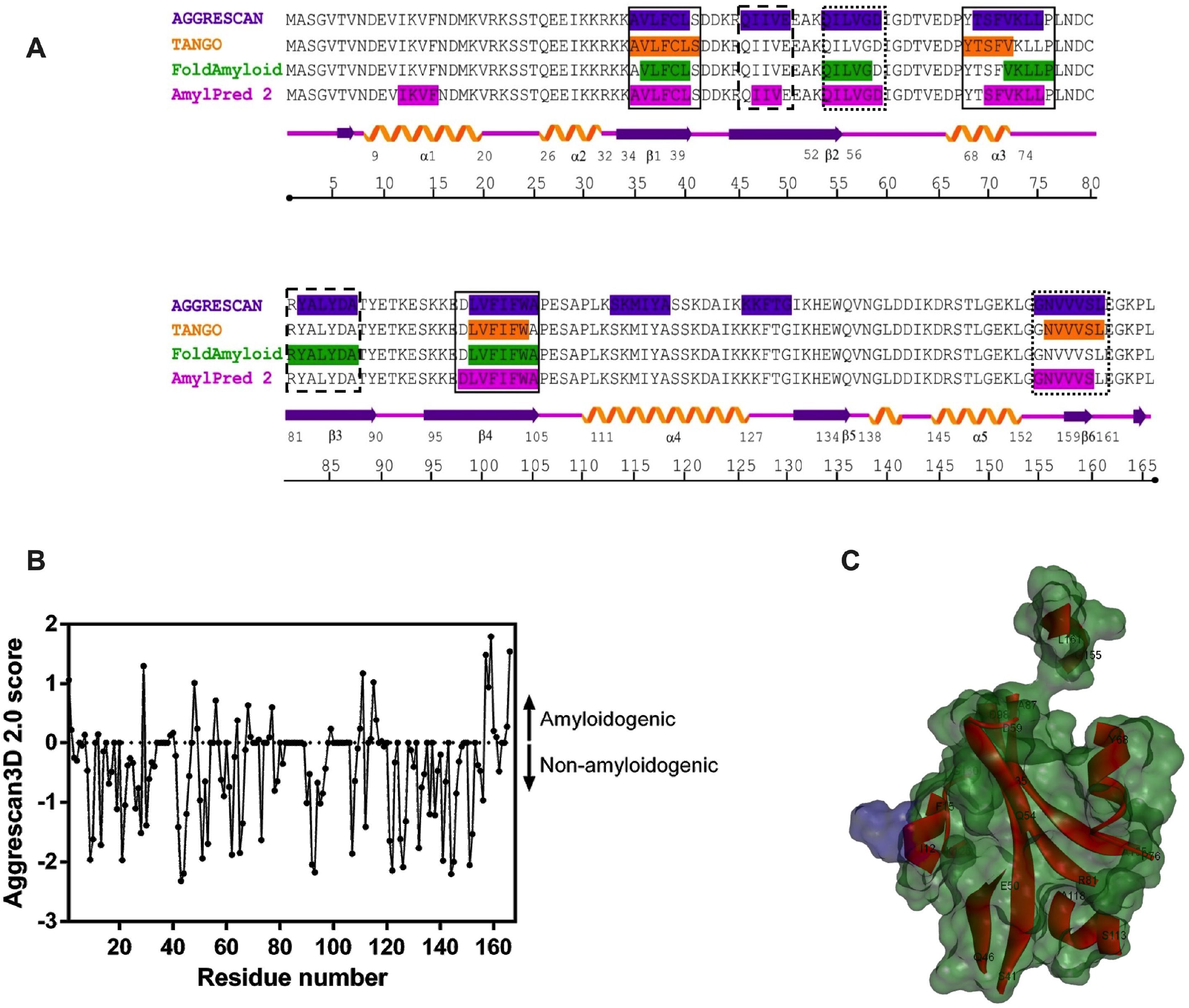
*In silico* analysis of cofilin-2 amyloid-forming tendency. The primary sequence and tertiary structural analyses of cofilin-2 were conducted for its amyloidogenic propensity using machine-learning-based open-access tools. (A) The image shows the aggregation-prone regions (APRs) in cofilin-2 primary sequence. The color codes; purple, orange, green, and magenta denote the APRs predicted by AGGRESCAN, TANGO, FoldAmyloid, and AmylPred 2, respectively. Dashed boxes: The APRs identified by at least two predictors, dotted boxes: predicted by at least three tools, and solid boxes: regions predicted by all the tools. (B) The graph shows the predictions by Aggrescan3D 2.0, which considers the residues with positive scores to be amyloidogenic. (C) The 3D cartoon of cofilin-2 with surface representation showing APRs as red ribbons and the surface denotes the extent of solvent accessibility of the regions, The regions under the blue surface are accessible to the solvent, and those under the green surface are buried inside the core.

We then analyzed the 3D structure of cofilin-2 by Aggrescan3D 2.0, which predicts the aggregation potential at per residue level in a folded protein structure. It predicted 31 residues in cofilin-2; among these, 10 were unique to Aggrescan3D 2.0, while the others were in common with those identified by the primary sequence analysis (Figure 1B and Table S2). Taking together, the primary and 3D structure analyses of cofilin-2 suggest that the protein holds high inherent amyloidogenic propensity. Further, we analyzed the solvent accessibility of the APRs predicted by primary and 3D prediction tools and found that all the APRs were buried inside the core. Of note, only K_13_ and I_29_ located in the predicted APRs were observed to be solvent-accessible (Figure 1C).

### 3.2. The surface exposure of APRs and their involvement in intermolecular interactions during cofilin-2 dimerization might induce its aggregation

We then asked whether the predicted APRs are involved in the protein-protein interactions during dimer formation through self-association. Therefore, we performed protein-protein rigid-body docking using the Z-Dock-3.0.2f server. The generated top 2000 poses were then re-ranked using the IRaPPA re-ranking algorithm [31] based on the energy parameters. The top-10 homodimers were selected for MD simulations, which revealed that 4 dimer complexes were stable under the given conditions. Later, we identified the residues falling at the dimer interface and involved in intermolecular interactions. We observed that most of the residues forming the intermolecular interactions, such as hydrogen bonding, salt bridge formation, electrostatic- and hydrophobic interactions, were in common with the predicted APRs of cofilin-2 (Figure 2, left panel). The details of the interacting residues are shown in Table S3. These data indicate that the APRs might be involved in cofilin-2 oligomerization.

**Figure 2:**
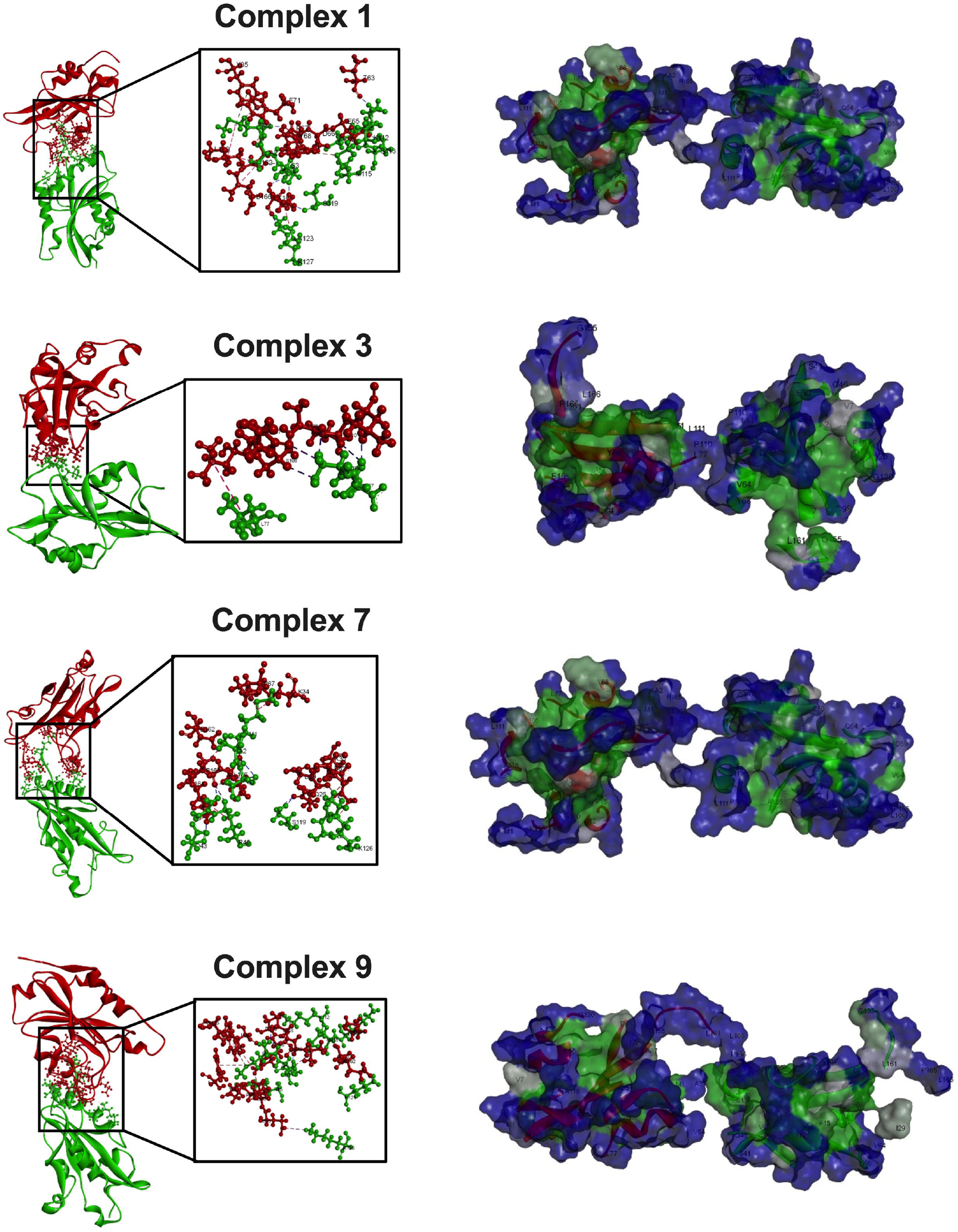
The APRs overlap with the interacting residues and become solvent-accessible in cofilin-2 homodimers. Cofilin-2 homodimers were generated and analyzed for their stability under given physicochemical conditions by molecular docking and MD simulations. (Left panel) The cartoon depictions of the stable cofilin-2 homodimer complexes show the residues of the APRs involved in intermolecular non-bond interactions. Chains A and B are shown in red and green, respectively. (Right panel) The surface models of cofilin-2 homodimers highlight the solvent accessibility of APRs. The solvent-accessible regions are shown in blue, and those which are buried are in green.

Further, we asked whether dimerization of cofilin-2 induces conformational changes and leads to exposure of APRs to the protein surface. Indeed, we found that the APRs which were buried in monomeric cofilin-2 (Figure 1C) get highly exposed to the surface and become solvent-accessible in cofilin-2 homodimers (Figure 2, right panel). The details of the solvent accessibility of the APRs in cofilin-2 homodimers are shown in Table S4. These data suggest that the surface exposure of APRs during dimerization of cofilin-2 might facilitate further interactions of cofilin-2 monomeric, dimeric or oligomeric species to form higher-order aggregates.

### 3.3. Cofilin-2 is capable of forming amyloid-like-fibrillar species *in vitro*

It remains unclear whether cofilin-2 forms amyloid-like structures under physiological conditions. We, therefore, explored the amyloid formation of cofilin-2 *in vitro*. The recombinant human cofilin-2 was incubated under physiological conditions for amyloid formation at different concentrations (25 μM and 50 μM). After 24 h of incubation, the amyloidogenic features of cofilin-2 aggregates were analyzed by employing native PAGE, CR binding assay, ThT binding assay, and fluorescence microscopy.

To identify the higher-order oligomers of cofilin-2, we performed native PAGE, in which various oligomeric species are separated based not only on the charge but also on the hydrodynamics of the protein. The native PAGE banding pattern showed both intermediate oligomers and mature aggregates, in addition to the monomeric cofilin-2 (Figure 3A).

**Figure 3:**
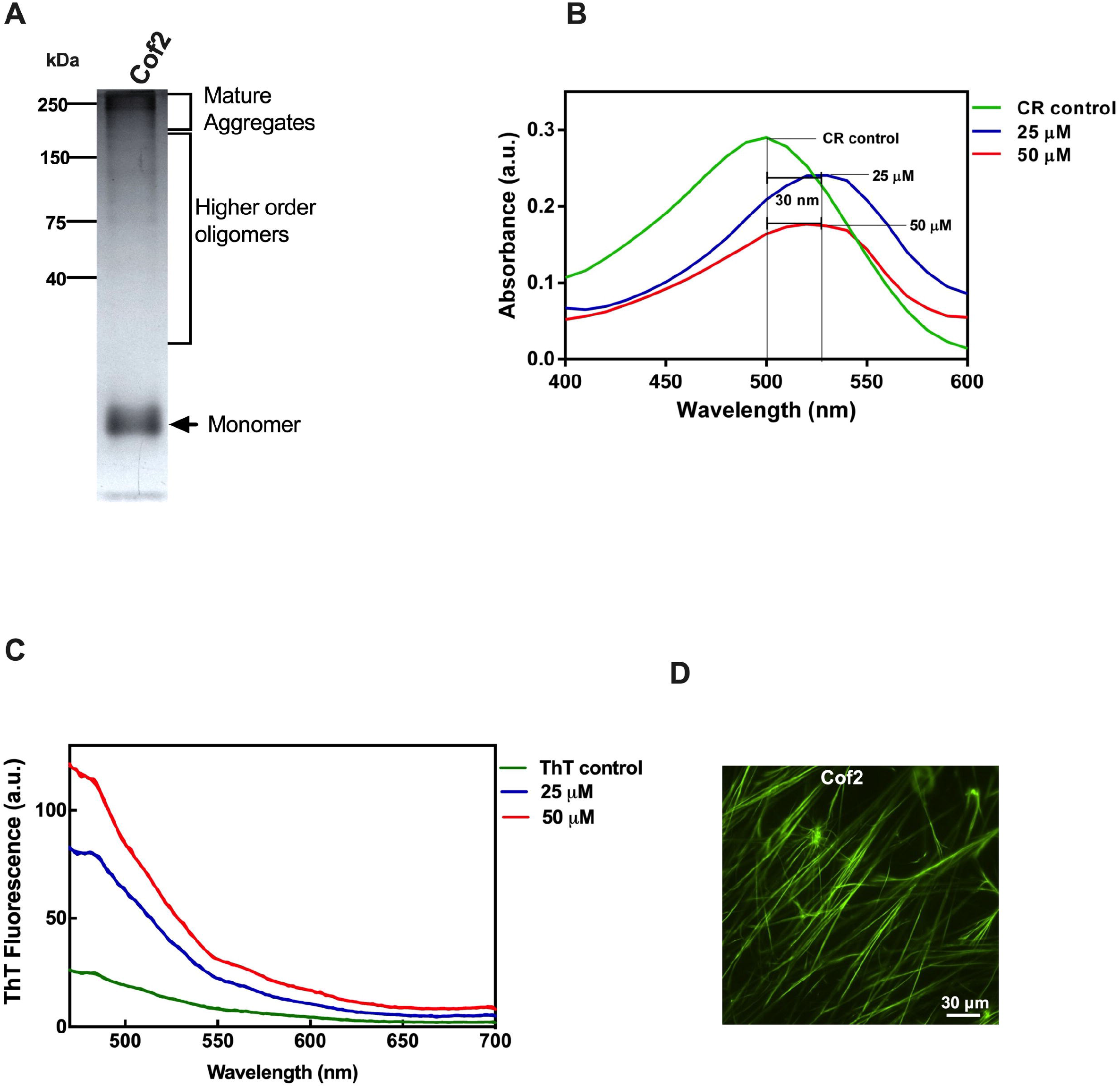
Cofilin-2 forms fibrillar amyloid-like structures *in vitro*. Cofilin-2 was assessed for its amyloid-like features under physiological conditions. (A) The native PAGE image shows the formation of a mixed population comprising mature aggregates, intermediate oligomers, and monomeric cofilin-2 at a concentration of 25 μM. (B) The absorption spectra of CR show that cofilin-2 aggregates exhibit a bathochromic shift of 30 nm in the absorption maxima of CR, signifying their amyloid-like characteristics. (C) The ThT fluorescence spectra denote that cofilin-2 aggregates show a drastic rise in the ThT fluorescence intensity, further forming cofilin-2 amyloid formation. (D) The micrograph shows that the ThT-stained cofilin-2 amyloids display long fibrillar morphology as visualized under a fluorescence microscope. Cof2: Cofilin-2.

We then performed the CR binding assay to identify the nature of these aggregates. It was observed that the aggregated cofilin-2 at 25 μM and 50 μM showed the maximum absorption at 530 nm signifying a 30 nm red shift from the CR control (500 nm), such a bathochromic shift is a characteristic of amyloids. (Figure 3B).

The ThT fluorescence assay further substantiated these data. The fluorescence intensity of ThT increased when bound to the aggregated cofilin-2 at concentrations of 25 μM and 50 μM by ∼3.5 fold and ∼5 fold, respectively, compared to the intensity of the unbound ThT (Figure 3C). These data suggest that cofilin-2 may form cross-β sheet-rich amyloid-like structures. Furthermore, we observed the ThT-stained cofilin-2 aggregates under a fluorescence microscope and found that cofilin-2 aggregates resemble long fibrillar morphology as observed for typical protein fibrils (Figure 3D). All these data strongly suggest that cofilin-2 has an intrinsic tendency to form amyloid-like structures under physiological conditions.

### 3.4. Cofilin-2 amyloid fibril seeds trigger the amyloidogenesis of globular proteins and amino acids

It has been recently shown that cofilin-1 promotes the aggregation of α-synuclein; therefore, we asked whether cofilin-2 also exhibits the potential to promote the aggregation of globular proteins. We co-incubated cofilin-2 amyloid seeds (5% w/w) with a mixed population of monomeric proteins (BSA, lysozyme, insulin, myoglobin, and cytochrome c). The cross-seeding effect of the preformed cofilin-2 amyloids was assessed by native PAGE, and it was found that the globular proteins, when cross-seeded with cofilin-2 amyloids, exhibited mixed species of higher-order oligomers and mature aggregates (Figure 4A). Further, the ThT fluorescence spectroscopy assay was performed, and we recorded a drastic increase in the fluorescence when the globular proteins were co-incubated with cofilin-2 amyloid seeds for 40 h (Figure 4B). The morphological characteristics of ThT-bound proteins’ aggregates seeded with cofilin-2 amyloids were assessed by fluorescence microscopy. We observed fibrillar morphology of aggregated proteins (Figure 4C).

**Figure 4:**
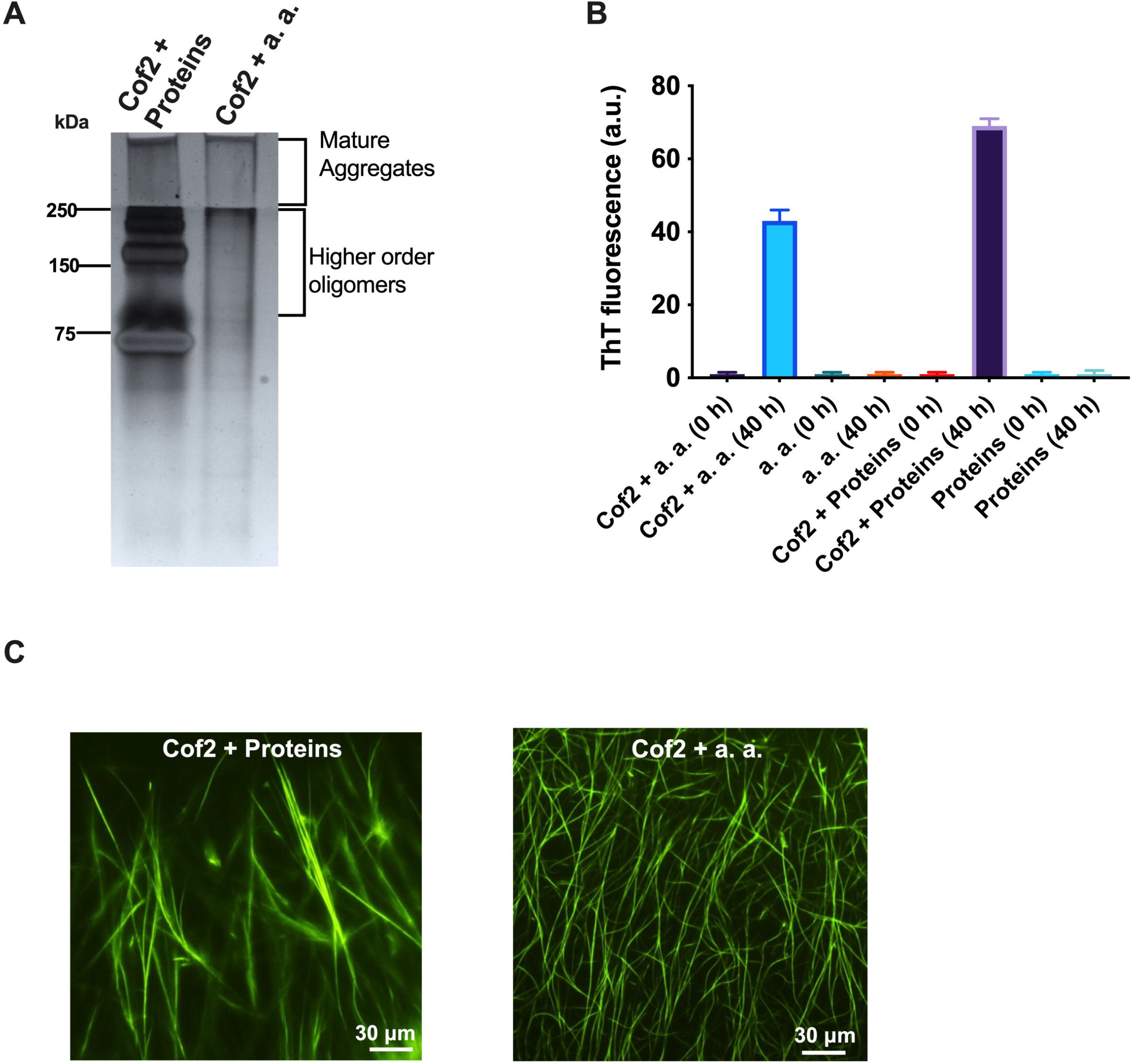
Cofilin-2 cross-seeds and facilitates the aggregation of globular proteins and amino acids. The effect of cofilin-2 on the aggregation of globular proteins and amino acids was assessed. (A) The non-denaturing gel shows that the globular proteins and amino acids seeded with 5% cofilin-2 amyloid seeds form higher-order oligomers and mature aggregates. (B) The bar diagram representing ThT fluorescence intensity at 490 nm shows that the species, which otherwise could not aggregate under physiological conditions, possess amyloidogenic characteristics upon cofilin-2 cross-seeding. (C) The fluorescence microscopy image shows that the proteins and amino acids cross-seeded with cofilin-2 exhibit fibrillar morphology. Cof2: Cofilin-2; a. a.: Amino acids.

Various studies suggest that not only globular proteins but also amino acids form amyloid-like structures in different disease conditions [31,32]. We spiked a mixture of amino acids (Phe, Tyr, Trp, Glu, Pro, and Ala) with cofilin-2 amyloid seeds, and amino acids’ aggregation was studied using native PAGE, ThT, and fluorescence microscopy. Indeed, we found that cofilin-2 amyloids were able to nucleate the aggregation of amino acids as well (Figure 4). These data suggest that cofilin-2 is capable of cross-seeding and promoting the amyloidogenesis of amino acids and globular proteins.

### 3.5. Both the cysteine residues (Cys-39 and Cys-80) in cofilin-2 are critical for its stability

Our recent study shows that Cys-80 in cofilin-1 is critical for its stability [20]. To analyze the effect of cysteine oxidation on cofilin-2 stability, we replaced the cysteine residues with aspartic acid (C to D mutation); this mimicked the formation of sulfonic acid, using Aggrescan3D 2.0. We analyzed the mutants based on the energy differences with respect to the WT structure. Any difference of ≥ 1 kcal/mol is indicative of a reduction in protein stability. The more the positive difference, the more unstable the structure is. We observed that the C39D and C80D mutants showed an energy difference of +1.75 kcal/mol and +2.17 kcal/mol, respectively, compared to the WT, and this difference further increased to +3.90 kcal/mol when the two cysteines were mutated simultaneously (Figure 5A). These data suggest that in contrast to cofilin-1, the oxidation of both cysteines in cofilin-2 synergistically impacts the protein stability

**Figure 5:**
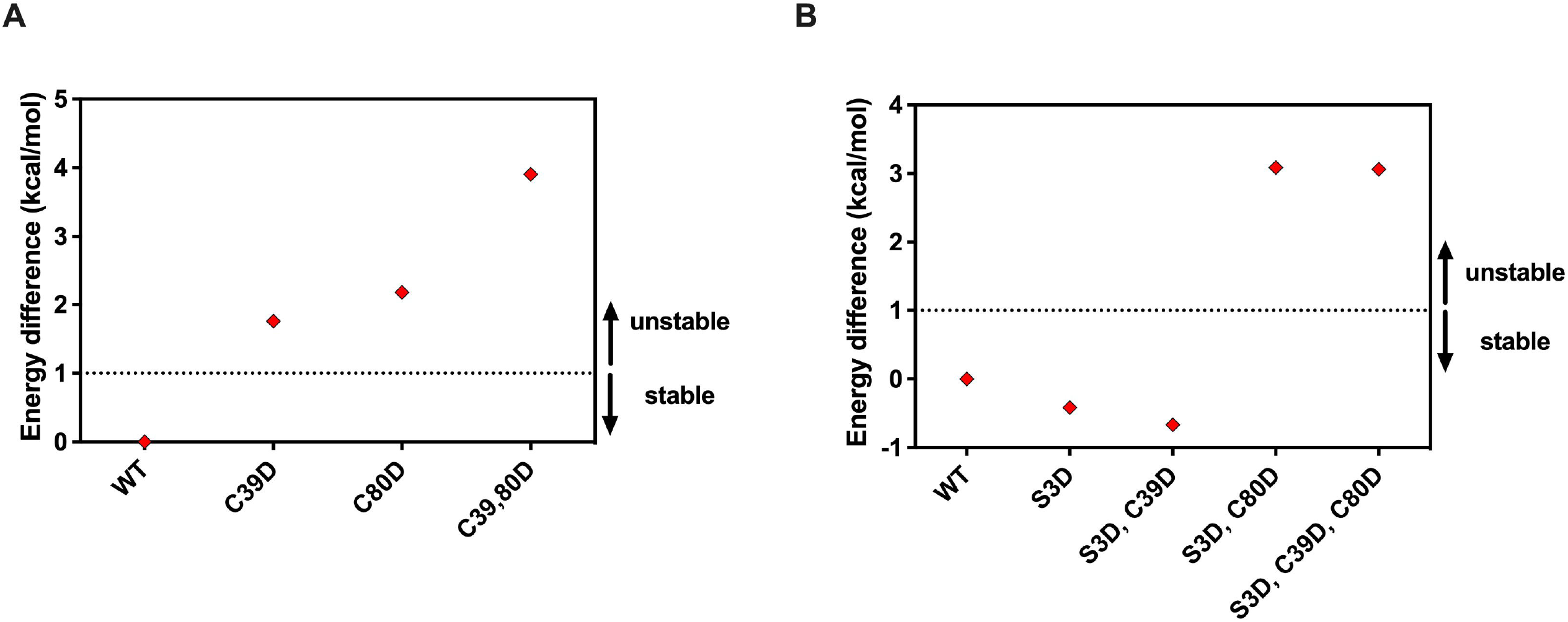
Both the cysteine residues (Cys-39 and Cys-80) are critical for cofilin-2 stability, while phosphorylation does not pose a significant impact. The thermodynamic stability of the oxidation and phosphorylation mimics (C to D and S to D mutants, respectively) was assessed by Aggrescan3D 2.0 server. (A) The graph denotes the energy differences caused by cysteine oxidation with respect to the WT. (B) The graph shows the energy differences of the phospho-mimic alone and in combination with cysteine oxidation with WT as the reference. Energy differences of ≥ 1 kcal/mol signify detrimental effects of the mutation on protein’s stability.

### 3.6. Phosphorylation did not significantly affect cofilin-2 stability

Previously, it was shown that phosphorylation of cofilin-2 promotes its aggregation under oxidative distress [4]. We asked whether the protein stability is affected by the phosphorylation of cofilin-2. The phosphomimetic mutant S3D alone tends to stabilize the protein but only to a minimal extent, with an energy difference of -0.41 kcal/mol. Interestingly, the S3D mutant reversed the detrimental effect of Cys-39 oxidation, stabilized the protein structure, and showed an energy difference of -0.66 kcal/mol. In contrast, the effect of oxidation at Cys-80 remained intact even after phosphorylation, as evident by the highly positive energy difference of +3.08, which is even higher than that observed for C80D mutation alone. This suggests that phosphorylation and Cys-80 oxidation cooperatively destabilize cofilin-2. Cofilin-2 phosphorylation with simultaneous oxidation both at Cys-39 and Cys-80 destabilizes the protein with an energy difference of +3.06 (Figure 5B). These data suggest that Cys-80 oxidation is more critical than Cys-39 oxidation in destabilizing cofilin-2 upon phosphorylation.

### 3.7. Cysteine oxidation causes partial structural unfolding in cofilin-2

The local structural perturbations due to oxidation partially unfold the protein and make it more prone to aggregation and amyloidogenesis [33]. We assessed the effect of cysteine oxidation on the structural features of cofilin-2 by *in silico* mutagenesis and MD simulations. The oxidation at Cys-39 or Cys-80 caused partial unfolding and reduced the helical and strand contents in cofilin-2 (Figure 6A and Figure S2). Moreover, C39D and C80D mutants showed a % SSE of 46.99% and 43.98%, respectively. This effect was further increased and was found to be 42.77% for the C39, 80D double oxidation mutant compared to the 59.04% in WT (Figure 6B). These reduced % SSE clarified that oxidation at Cys-39 or Cys-80 in cofilin-2 resulted in partial unfolding in the native protein.

**Figure 6:**
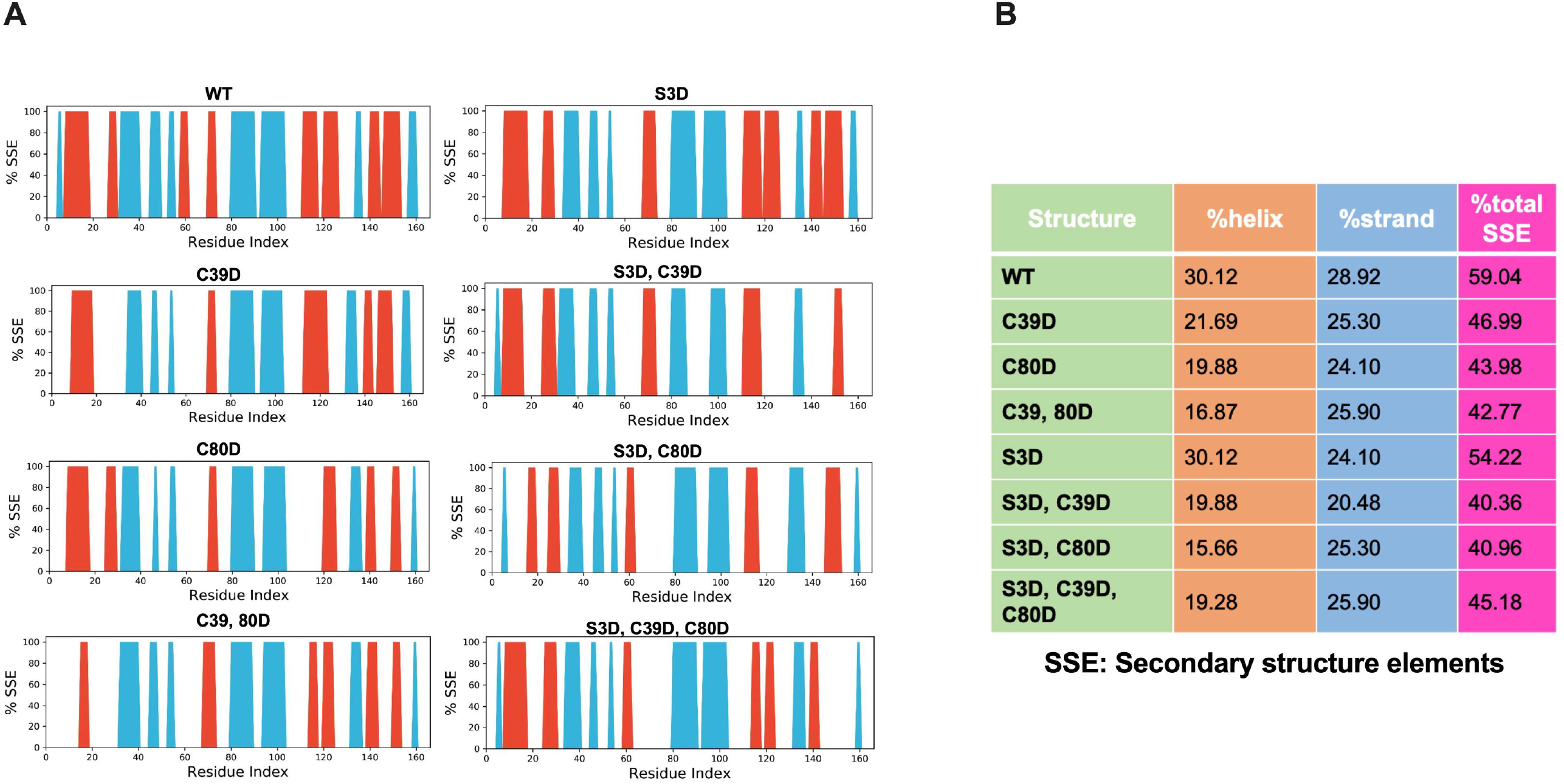
Cysteine oxidation in cofilin-2 partially unfolds the protein. The cysteine oxidation and phospho-mimics were assessed for their impact on the structural characteristics of cofilin-2 by *in silico* mutagenesis and MD simulations. (A) The graphs denote the alteration in the helical (orange) and strand (cyan) content of cofilin-2 upon cysteine oxidation and phosphorylation of cofilin-2. (B) The table shows the detailed scoring of the secondary structure content of WT and the mutants.

We also analyzed the effect of phosphorylation on the structural characteristics of cofilin-2 and found that S3D alone showed a % SSE of 54.22%, which is close to that observed for the WT. Further, the phosphorylation mimic of cofilin-2 partially enhances the unfolding caused by Cys-39 or Cys-80 oxidation, with % SSE values of 40.36% and 40.96%, respectively (Figure 6A, Figure 6B and Figure S2). Interestingly, phosphorylation of cofilin-2 slightly reversed the unfolding effect caused by simultaneous oxidation of Cys-39 and Cys-80. These data suggest that phosphorylation did not significantly affect the protein structure in relation to cofilin-2 aggregation

## 4. Discussion

The available literature suggests that cofilin-2, the muscle isoform of the ADF/cofilin family, is involved in the etiology of various pathological conditions, such as protein aggregate accumulations in cardiomyopathies [17]. Pignataro et al. suggested that the phosphorylated cofilin-2 might form amyloid-like fibrillar structures, but only under pathophysiological conditions, such as oxidative distress [4]. However, the cofilin-2 amyloid formation under physiological conditions remains unexplored. The current study shows that, like the non-muscle form, the muscle cofilin also exhibits the innate tendency to form amyloids, and cysteine oxidation might accelerate cofilin-2 amyloidogenesis. It also suggests that the enhanced solvent-accessibility of APRs during the early stages of aggregation might pose a mechanistic approach towards cofilin-2 amyloidogenesis. Further, the potential of cofilin-2 amyloids to induce the aggregation of other proteinaceous species might aggravate the pathophysiology of hetero-aggregation linked diseases.

Compared to cofilin-1, cofilin-2 possesses more APRs as identified by the primary and tertiary structural analyses. The primary sequence analysis deciphered 9 and 10 APRs in cofilin-1 and -2, respectively, while the tertiary structure analysis highlighted 27 residues in cofilin-1 (Table S5) and 31 residues in cofilin-2 to be prone to aggregation. Moreover, the *in vitro* experiments suggested that cofilin-2 gets aggregated to form amyloid-like structures at a concentration of 25 μM, while cofilin-1 was observed to form similar structures at ∼3 fold higher concentrations *in vitro*. Cofilin-2 formed longer amyloid fibrils compared to the shorter fibrils of cofilin-1 [20]. These differences in the number of aggregation-prone residues/regions, the concentration required to form aggregates, and the morphology clearly indicate that cofilin-2 is more prone to amyloidogenesis. Our current study hints towards the possibility that the local cofilin-2 concentration regulates its oligomerization similar to cofilin-1 and that the increase in the local concentration above the threshold under pathophysiological conditions promotes its aggregation leading to amyloid-like fibrillar structure formation. We cannot rule out that cofilin-2, like cofilin-1, forms small oligomers, such as dimers or tetramers, regulating its physiological functions, such as actin dynamization.

It was reported that the aggregation-prone regions of an amyloidogenic protein overlap with protein interfaces and interaction sites, indicating a competition between the formation of functional complexes under physiological state and aggregates under pathophysiological conditions [34]. Here, we employed protein-protein docking and MD simulations to generate cofilin-2 homodimers. It is interesting to note that most of the interacting residues overlapped with the identified aggregation-prone regions/residues, indicating that the APRs are critical players in the initial association of cofilin-2 monomers and dimer formation. Moreover, the buried APRs get exposed in these homodimers, implying mechanistic insight that the exposure of APRs to the surface and their increased solvent accessibility may promote cofilin-2 aggregation and eventually amyloid formation.

The hallmarks of the amyloid-linked diseases, such as plaques, tangles, Hirano bodies, Lewy bodies, are heterogeneous and contain a myriad of protein aggregates, establishing a cross-talk and contributing to the etiology of the associated diseases [35]. The phenomenon of cross-seeding wherein preformed amyloids act as seeds and eliminate the requirement for primary nucleation poses an underlying explanation to the biologically relevant hetero-aggregation of proteinaceous species [36]. Moreover, a recent report showed that cofilin-1 facilitates the aggregation of α-synuclein, leading to neuronal dysfunction, indicating a possible link of cofilin-1 to Parkinson’s disease [37]. In the present study, we propose that cofilin-2 amyloids might promote the template-dependent aggregation of other proteins and amino acids in the cellular milieu by providing seeds and facilitating the rate-limiting nucleation stage during amyloid formation. Indeed, cofilin-2 amyloid seeds were able promote amyloid-formation in various globular proteins and amino acids, which otherwise do not form such structures under the given conditions.

Oxidation of cellular proteins tends to thermodynamically destabilize the proteins, leading to the formation of aggregates, loss of function, and cytotoxicity [33]. Protein aggregation is sensitive to oxidation, in fact so much that oxidation at even a single cysteine residue is sufficient to alter the energy landscape and kinetic stability of a protein, thereby taking it towards the amyloid formation pathway [38]. Several reports suggest that cofilins are potential targets for oxidation at their cysteine residues, resulting in disulfide bonding and sulfonic acid formation. These perturbations cause structural and functional variations in the protein, which further leads to the formation of toxic actin-cofilin rods, known to be involved in neuronal damage [13,39]. The present study shows that in contrast to cofilin-1, oxidation at either of the two conserved cysteines (Cys-39 and Cys-80) destabilizes cofilin-2 [20]. This effect is further complemented by the simultaneous oxidation of the two residues. Therefore, we propose that cofilin-2 is more susceptible to the changes caused by oxidation than cofilin-1, which, however, possesses two additional cysteines but only one, i.e., Cys-80 is critical for its stability. A recent study reported that both phosphorylation and oxidation of cofilin-2 are essential for amyloid-like fibrils’ formation [4]. In contrast, our *in silico* and *in vitro* data show that phosphorylation is not required for amyloid formation. The possible explanation could be that a very low concentration of cofilin-2 (5 μM) was used in the previous study, at which the protein aggregation might not be possible in its unmodified form.

The oxidized forms of proteins attain partially unfolded tertiary structures and readily form aggregates. These oxidized proteins might act as intermediate conformers in the amyloid fibril formation [33]. Our mutagenesis and MD simulation data give a deeper insight into the structural changes occurring due to oxidation. It suggests that both oxidized cysteines reduce the total folded content and compactness of the protein. The transition of helical and strand regions into loops indicates localized unfolding, an initial step in the amyloid formation. Therefore, it is clear that cysteines’ oxidation in cofilin-2 facilitates its aggregation by destabilization and loss of native structure.

Taking together, we could successfully describe that cofilin-2 possesses an innate tendency to form amyloid-like structures under physiological conditions, which is facilitated by the high solvent-accessibility of APRs during the early stages of aggregation, i.e., dimerization. Cofilin fibrillation might negatively impact the homeostasis by altering its actin dynamizing property and affecting its physiological functions, such as cell migration. Further, these amyloids may pose cytotoxic effects due to the possible gain of toxic function. The cofilin-2 amyloids hold the potential to cross-seed both globular proteins and amino acids, thereby aggravating the etiology of amyloid-linked diseases involving hetero-aggregation of proteinaceous species. These results may provide foundational knowledge for the molecular origin of diseased states, such as myopathies. Therefore, we speculate that the accumulation of cofilin-2 into amyloids could be a novel target for the diagnosis and therapeutics of cytoskeletal deformities in conformational disorders.

## Authors’ contribution

**Vibha Kaushik**: experimentation, analysis, visualization, writing, reviewing, and editing. **Eva-Maria Hanschmann**: cloning, reviewing, and editing. **Daniela Brünnert**: visualization, reviewing, and editing. **Kumari Prerna**: spectroscopy experiments. **Bibin G. Anand**: cross-seeding experiments. **Phulwanti Kumari Sharma**: manuscript reviewing. **Karunakar Kar**: resources, reviewing, and editing. **Pankaj Goyal**: conceptualization, supervision, resources, analysis, visualizations, reviewing, and editing.

## Acknowledgments

V.K. and P.K.S. are recipients of fellowships from the University Grants Commission (JRF-21/06/2015 (i) EU-V/2061530804 to V.K and JRF-20/12/2015 (ii) EU-V/2121530832 to P.K.S.), Government of India. We thank the Department of Chemistry, Central University of Rajasthan, for providing the fluorimeter facility. We are also grateful to Dr. Akhil Agrawal, Department of Microbiology, Central University of Rajasthan, and Dr. Suneel Kateriya, School of Biotechnology, Jawaharlal Nehru University, New Delhi, for their support.

## Supplementary figures

**Figure S1:** The cofilin-2 protein was modeled using the I-TASSER server, and the quality was then validated by PSVS (A) The cartoon representation of the best 3D model of cofilin-2 selected for the study. Cyan: β-sheets, red: α-helices, green: turns, and white: coils. (B) The Ramachandran plot of the modeled cofilin-2 structure.

**Figure S2:** The 3D cartoon depictions of cofilin-2 (WT) and its mutants showing the alterations in the secondary structure caused by oxidation and phosphorylation mimics.

**Table S1:** The table shows the APRs as predicted by (A) AGGRESCAN, (B) TANGO, and (C) FoldAmyloid with their respective scores for each residue of the APRs.

**Table S2:** The table includes the APRs and their respective scores in cofilin-2 as predicted by the Aggrescan3D 2.0 server.

**Table S3:** The interacting residues in cofilin-2 stable homodimer complexes (A) Complex 1, (B) Complex 3, (C) Complex 7, (D) Complex 9. The residues highlighted in yellow are APRs overlapping the interacting residues.

**Table S4:** The table denotes solvent accessibility of APRs in stable cofilin-2 homodimers (A) Complex 1, (B) Complex 3, (C) Complex 7, and (D) Complex 9. Blue, Cyan, and Green represent % residue solvent accessibility of more than 25%, between 10% to 25%, and less than 10%, respectively.

**Table S5:** The table includes the APRs and their respective scores in cofilin-2 as predicted by the Aggrescan3D 2.0 server.

