## Supplementary info IS1 for "Computational and biochemical analyses reveal that cofilin-2 self assembles into amyloid-like structures and promotes the aggregation of other proteinaceous species: Pathogenic relevance to myopathies"

**Supplementary information**

**SI-1: Homology modeling and model validation**

The crystal structure of human cofilin-2 is not available, and thus, we developed a 3D model of the structure by Iterative-Threading ASSEmbly Refinement (I-TASSER) server [1]. It exploits the multiple threading approach to identify structural templates and generates a full-length atomic model by iterative template-based fragment assembly simulations. Using this method, we generated 4 models and selected the best-modeled structure (Figure S1A) based on the highest confidence score (C-score) (0.22), which denotes the convergence parameters of the structure and the significance of threading template alignments and assembly simulations. The selected model structure was then subjected to molecular dynamics simulations for 10 ns using the Desmond molecular dynamics module (Schrӧdinger Inc., USA) [2] to remove the steric clashes and obtain a stable structure. Later, the model was validated by Protein Structure Validation Suite (PSVS) [3], which includes various algorithms for model quality assessment. It suggested that 99.4% of the regions in the selected model lay in the favored, allowed, and generously allowed areas of the Ramachandran plot as per the analysis with Procheck, and 98.1% region of the model was favored according to Richardson Lab's Molprobity algorithm (Figure S1B). All the algorithms incorporated in the PSVS suite assigned a Z-score ranging from -4.01 to +0.69 to the model. Thus, the modeled structure of cofilin-2 could be considered a good model for further analysis in the study.
