## Supplementary material for "Computational and biochemical analyses reveal that cofilin-2 self assembles into amyloid-like structures and promotes the aggregation of other proteinaceous species: Pathogenic relevance to myopathies": Table S1-S5

**Table S1:** The table shows the APRs as predicted by (A) AGGRESCAN, (B) TANGO, and (C) FoldAmyloid with their respective scores for each residue of the APRs.

A. **AGGRESCAN**

| **Predicted APRs** | **Residue number** | **Residue name** | **Average of aggregation propensity values per amino acid**  **(a^4^v)** | **Hot spot area (HSA)** | **Normalized hot spot area per 100 residue hot spot**  **(NHSA)** | **Average a4v in each hot spot**  **(a^4^vAHS)** |
| --- | --- | --- | --- | --- | --- | --- |
| **A35-L40** | 35 | A | 0.227 | 3.389 | 0.565 | 0.545 |
|  | 36 | V | 0.491 | 3.389 | 0.565 | 0.545 |
|  | 37 | L | 0.821 | 3.389 | 0.565 | 0.545 |
|  | 38 | F | 0.912 | 3.389 | 0.565 | 0.545 |
|  | 39 | C | 0.655 | 3.389 | 0.565 | 0.545 |
|  | 40 | L | 0.165 | 3.389 | 0.565 | 0.545 |
| **Q46-E50** | 46 | Q | 0 | 0.523 | 0.105 | 0.085 |
|  | 47 | I | 0.061 | 0.523 | 0.105 | 0.085 |
|  | 48 | I | -0.008 | 0.523 | 0.105 | 0.085 |
|  | 49 | V | 0.164 | 0.523 | 0.105 | 0.085 |
|  | 50 | E | 0.207 | 0.523 | 0.105 | 0.085 |
| **Q54-D59** | 54 | Q | 0.169 | 1.591 | 0.265 | 0.245 |
|  | 55 | I | 0.295 | 1.591 | 0.265 | 0.245 |
|  | 56 | L | 0.038 | 1.591 | 0.265 | 0.245 |
|  | 57 | V | 0.431 | 1.591 | 0.265 | 0.245 |
|  | 58 | G | 0.53 | 1.591 | 0.265 | 0.245 |
|  | 59 | D | 0.008 | 1.591 | 0.265 | 0.245 |
| **A69-M74** | 69 | T | 0.269 | 4.117 | 0.588 | 0.568 |
|  | 70 | S | 0.398 | 4.117 | 0.588 | 0.568 |
|  | 71 | F | 0.643 | 4.117 | 0.588 | 0.568 |
|  | 72 | V | 0.675 | 4.117 | 0.588 | 0.568 |
|  | 73 | K | 0.65 | 4.117 | 0.588 | 0.568 |
|  | 74 | L | 0.889 | 4.117 | 0.588 | 0.568 |
|  | 75 | L | 0.452 | 4.117 | 0.588 | 0.568 |

| **Y82-A87** | 82 | Y | 0.17 | 1.041 | 0.173 | 0.154 |
| --- | --- | --- | --- | --- | --- | --- |
|  | 83 | A | 0.17 | 1.041 | 0.173 | 0.154 |
|  | 84 | L | 0.079 | 1.041 | 0.173 | 0.154 |
|  | 85 | Y | 0.233 | 1.041 | 0.173 | 0.154 |
|  | 86 | D | 0.233 | 1.041 | 0.173 | 0.154 |
|  | 87 | A | 0.036 | 1.041 | 0.173 | 0.154 |
| **L99-A105** | 99 | L | 0.339 | 5.704 | 0.815 | 0.795 |
|  | 100 | V | 0.722 | 5.704 | 0.815 | 0.795 |
|  | 101 | F | 1.072 | 5.704 | 0.815 | 0.795 |
|  | 102 | I | 1.329 | 5.704 | 0.815 | 0.795 |
|  | 103 | F | 1.084 | 5.704 | 0.815 | 0.795 |
|  | 104 | W | 0.655 | 5.704 | 0.815 | 0.795 |
|  | 105 | A | 0.362 | 5.704 | 0.815 | 0.795 |
| **S113-A118** | 113 | S | 0.232 | 2.041 | 0.34 | 0.32 |
|  | 114 | K | 0.445 | 2.041 | 0.34 | 0.32 |
|  | 115 | M | 0.243 | 2.041 | 0.34 | 0.32 |
|  | 116 | I | 0.334 | 2.041 | 0.34 | 0.32 |
|  | 117 | Y | 0.334 | 2.041 | 0.34 | 0.32 |
|  | 118 | A | 0.334 | 2.041 | 0.34 | 0.32 |
| **K126-L130** | 126 | K | 0.084 | 0.22 | 0.044 | 0.024 |
|  | 127 | K | 0.013 | 0.22 | 0.044 | 0.024 |
|  | 128 | F | 0.013 | 0.22 | 0.044 | 0.024 |
|  | 129 | T | 0.013 | 0.22 | 0.044 | 0.024 |
|  | 130 | G | -0.002 | 0.22 | 0.044 | 0.024 |
| **G155-L161** | 155 | G | 0.181 | 2.95 | 0.421 | 0.401 |
|  | 156 | N | 0.541 | 2.95 | 0.421 | 0.401 |
|  | 157 | V | 0.302 | 2.95 | 0.421 | 0.401 |
|  | 158 | V | 0.576 | 2.95 | 0.421 | 0.401 |
|  | 159 | V | 0.451 | 2.95 | 0.421 | 0.401 |
|  | 160 | S | 0.56 | 2.95 | 0.421 | 0.401 |
|  | 161 | L | 0.199 | 2.95 | 0.421 | 0.401 |

B. **TANGO**

| **Predicted APRs** | **Residue number** | **Residue name** | **Beta sheet** | **Beta turn** | **Alpha helix** | **Beta aggregation** | **Helix aggregation** |
| --- | --- | --- | --- | --- | --- | --- | --- |
| **A35-S41** | 35 | A | 1.2 | 0.1 | 0.505 | 21.675 | 0.006 |
|  | 36 | V | 2.8 | 0 | 0.277 | 59.863 | 0.016 |
|  | 37 | L | 4.4 | 0 | 0.277 | 59.863 | 0.016 |
|  | 38 | F | 6.1 | 0 | 0.15 | 59.863 | 0.016 |
|  | 39 | C | 6.1 | 0 | 0 | 59.863 | 0.016 |
|  | 40 | L | 4.3 | 0.3 | 0 | 59.443 | 0.016 |
|  | 41 | S | 1.8 | 1.4 | 0.656 | 10.651 | 0.003 |
| **Y68-V72** | 68 | Y | 3.1 | 0.3 | 2.003 | 4.083 | 0 |
|  | 69 | T | 4.6 | 0.3 | 2.003 | 4.083 | 0 |
|  | 70 | S | 5.6 | 0 | 2.003 | 4.083 | 0 |
|  | 71 | F | 4.8 | 0 | 2.003 | 4.083 | 0 |
|  | 72 | V | 10.3 | 0 | 1.395 | 4.083 | 0 |
| **L99-W104** | 99 | L | 1.6 | 0.5 | 0.176 | 71 | 0.407 |
|  | 100 | V | 4.5 | 0 | 0.176 | 98.091 | 0.563 |
|  | 101 | F | 5 | 0 | 0.176 | 98.123 | 0.563 |
|  | 102 | I | 4.6 | 0 | 0.176 | 98.123 | 0.563 |
|  | 103 | F | 4.7 | 0 | 0.176 | 98.123 | 0.563 |
|  | 104 | W | 4 | 0 | 0 | 83.694 | 0.48 |
| **N156-L161** | 156 | N | 2 | 0.5 | 0 | 1.276 | 0 |
|  | 157 | V | 14.1 | 0.2 | 0 | 19.184 | 0.003 |
|  | 158 | V | 18.4 | 0 | 0 | 19.184 | 0.003 |
|  | 159 | V | 20.9 | 0 | 0 | 19.184 | 0.003 |
|  | 160 | S | 10 | 0 | 0 | 18.698 | 0.003 |
|  | 161 | L | 5.5 | 0.4 | 0 | 18.567 | 0.002 |

C. **FoldAmyloid**

| **Predicted APRs** | **Residue number** | **Residue name** | **Fold value** |
| --- | --- | --- | --- |
| **V36-L40** | 36 | V | 22.806 |
|  | 37 | L | 23.976 |
|  | 38 | F | 25.070 |
|  | 39 | C | 23.922 |
|  | 40 | L | 22.332 |
| **Q54-G58** | 54 | Q | 21.572 |
|  | 55 | I | 22.380 |
|  | 56 | L | 22.268 |
|  | 57 | V | 21.904 |
|  | 58 | G | 21.904 |
| **V72-P76** | 72 | V | 22.466 |
|  | 73 | K | 23.900 |
|  | 74 | L | 21.950 |
|  | 75 | L | 22.236 |
|  | 76 | P | 22.400 |
| **R81A87** | 81 | R | 21.556 |
|  | 82 | Y | 23.146 |
|  | 83 | A | 23.628 |
|  | 84 | L | 22.904 |
|  | 85 | Y | 21.696 |
|  | 86 | D | 21.680 |
|  | 87 | A | 21.794 |
| **L99-A105** | 99 | L | 22.268 |
|  | 100 | V | 23.918 |
|  | 101 | F | 25.872 |
|  | 102 | I | 26.496 |
|  | 103 | F | 25.688 |
|  | 104 | W | 23.738 |
|  | 105 | A | 22.088 |

**Table S2:** The table includes the APRs and their respective scores in cofilin-2 as predicted by the Aggrescan3D 2.0 server.

| **Residue number** | **Residue name** | **Aggrescan3D 2.0 score** |
| --- | --- | --- |
| 1 | M | 1.0646 |
| 2 | A | 0.2187 |
| 7 | V | 0.1372 |
| 12 | I | 0.1417 |
| 29 | I | 1.2961 |
| 39 | C | 0.1270 |
| 40 | L | 0.1676 |
| 48 | I | 1.0154 |
| 49 | V | 0.2415 |
| 56 | L | 0.7207 |
| 64 | V | 0.3820 |
| 68 | Y | 0.6384 |
| 69 | T | 0.1028 |
| 72 | V | 0.0575 |
| 76 | P | 0.0930 |
| 77 | L | 0.6076 |
| 99 | L | 0.2335 |
| 105 | A | 0.0072 |
| 110 | P | 0.2385 |
| 111 | L | 1.1773 |
| 114 | K | 0.0681 |
| 115 | M | 1.0272 |
| 116 | I | 0.3915 |
| 118 | A | 0.0272 |
| 157 | V | 1.4872 |
| 158 | V | 0.9417 |
| 159 | V | 1.7949 |
| 160 | S | 0.2010 |
| 161 | L | 0.1000 |
| 165 | P | 0.2737 |
| 166 | L | 1.5430 |

**Table S3:** The interacting residues in cofilin-2 stable homodimer complexes (A) Complex 1, (B) Complex 3, (C) Complex 7, (D) Complex 9. The residues highlighted in yellow are APRs overlapping the interacting residues.

A. **Complex 1**

| **From** | **To** | **From Chemistry** | **To Chemistry** | **Interaction** | **Type** | **Distance** |
| --- | --- | --- | --- | --- | --- | --- |
| B:MET1:H2 | A:GLU162:OE2 | H-Donor;Positive | H-Acceptor;Negative | B:MET1:H2 - A:GLU162:OE2 | Hydrogen Bond;Electrostatic; Salt Bridge;Attractive Charge | 1.80181 |
| B:LYS44:HZ2 | A:GLU65:OE2 | H-Donor;Positive | H-Acceptor;Negative | B:LYS44:HZ2 - A:GLU65:OE2 | Hydrogen Bond;Electrostatic; Salt Bridge;Attractive Charge | 1.56895 |
| B:LYS112:HZ1 | A:GLU65:OE1 | H-Donor;Positive | H-Acceptor;Negative | B:LYS112:HZ1 - A:GLU65:OE1 | Hydrogen Bond;Electrostatic; Salt Bridge;Attractive Charge | 1.56272 |
| A:LYS164:H | B:GLY4:O | H-Donor | H-Acceptor | A:LYS164:H - B:GLY4:O | Conventional Hydrogen Bond | 2.72974 |
| A:LYS164:HZ1 | B:SER3:OG | H-Donor | H-Acceptor | A:LYS164:HZ1 - B:SER3:OG | Conventional Hydrogen Bond | 1.73066 |
| A:LYS164:HZ3 | B:SER119:O | H-Donor | H-Acceptor | A:LYS164:HZ3 - B:SER119:O | Conventional Hydrogen Bond | 1.64381 |
| B:MET1:H1 | A:PRO67:O | H-Donor | H-Acceptor | B:MET1:H1 - A:PRO67:O | Conventional Hydrogen Bond | 2.00434 |
| B:MET1:H3 | A:TYR85:OH | H-Donor | H-Acceptor | B:MET1:H3 - A:TYR85:OH | Conventional Hydrogen Bond | 2.23996 |
| B:ALA2:H | A:GLU162:OE2 | H-Donor | H-Acceptor | B:ALA2:H - A:GLU162:OE2 | Conventional Hydrogen Bond | 1.70707 |
| B:LYS112:HZ2 | A:THR63:O | H-Donor | H-Acceptor | B:LYS112:HZ2 - A:THR63:O | Conventional Hydrogen Bond | 1.85294 |
| B:SER113:HG | A:GLU65:OE2 | H-Donor | H-Acceptor | B:SER113:HG - A:GLU65:OE2 | Conventional Hydrogen Bond | 1.88915 |
| B:LYS127:HZ1 | A:LYS164:O | H-Donor | H-Acceptor | B:LYS127:HZ1 - A:LYS164:O | Conventional Hydrogen Bond | 2.45638 |
| B:SER3:HA | A:ASP66:OD2 | H-Donor | H-Acceptor | B:SER3:HA - A:ASP66:OD2 | Carbon Hydrogen Bond | 2.69537 |
| B:GLY4:HA2 | A:GLU162:O | H-Donor | H-Acceptor | B:GLY4:HA2 - A:GLU162:O | Carbon Hydrogen Bond | 2.78536 |
| B:MET115:SD | A:TYR68 | Sulfur | Pi-Orbitals | B:MET115:SD - A:TYR68 | Pi-Sulfur | 5.22362 |
| A:LEU149 | B:MET1 | Alkyl | Alkyl | A:LEU149 - B:MET1 | Alkyl | 5.28159 |
| B:ALA2 | A:LEU149 | Alkyl | Alkyl | B:ALA2 - A:LEU149 | Alkyl | 5.21431 |
| B:ALA2 | A:LEU166 | Alkyl | Alkyl | B:ALA2 - A:LEU166 | Alkyl | 4.16694 |
| B:ALA123 | A:LYS164 | Alkyl | Alkyl | B:ALA123 - A:LYS164 | Alkyl | 4.13767 |
| A:PHE71 | B:MET1 | Pi-Orbitals | Alkyl | A:PHE71 - B:MET1 | Pi-Alkyl | 5.21719 |
| A:TYR85 | B:MET1 | Pi-Orbitals | Alkyl | A:TYR85 - B:MET1 | Pi-Alkyl | 4.83551 |

B. **Complex 3**

| **From** | **To** | **From Chemistry** | **To Chemistry** | **Interaction** | **Type** | **Distance** |
| --- | --- | --- | --- | --- | --- | --- |
| A:LEU111:H | B:GLU107:O | H-Donor | H-Acceptor | A:LEU111:H - B:GLU107:O | Conventional Hydrogen Bond | 2.60325 |
| A:SER108:HA | B:SER108:OG | H-Donor | H-Acceptor | A:SER108:HA - B:SER108:OG | Carbon Hydrogen Bond | 2.64974 |
| A:PRO110:HA | B:GLU107:O | H-Donor | H-Acceptor | A:PRO110:HA - B:GLU107:O | Carbon Hydrogen Bond | 2.78936 |
| B:SER108:HB1 | A:GLU107:O | H-Donor | H-Acceptor | B:SER108:HB1 - A:GLU107:O | Carbon Hydrogen Bond | 2.94008 |
| B:SER108:HB2 | A:ALA109:O | H-Donor | H-Acceptor | B:SER108:HB2 - A:ALA109:O | Carbon Hydrogen Bond | 2.52786 |
| A:LEU77 | B:LEU77 | Alkyl | Alkyl | A:LEU77 - B:LEU77 | Hydrophobic Alkyl | 5.47464 |

C. **Complex 7**

| **From** | **To** | **From Chemistry** | **To Chemistry** | **Interaction** | **Type** | **Distance** |
| --- | --- | --- | --- | --- | --- | --- |
| A:LYS30:HZ2 | B:ASP122:OD1 | H-Donor;Positive | H-Acceptor;Negative | A:LYS30:HZ2 - B:ASP122:OD1 | Hydrogen Bond;Electrostatic, Salt Bridge;Attractive Charge | 1.74014 |
| B:MET1:H3 | A:GLU162:OE2 | H-Donor;Positive | H-Acceptor;Negative | B:MET1:H3 - A:GLU162:OE2 | Hydrogen Bond;Electrostatic, Salt Bridge;Attractive Charge | 1.57143 |
| B:LYS126:HZ3 | A:ASP59:OD2 | H-Donor;Positive | H-Acceptor;Negative | B:LYS126:HZ3 - A:ASP59:OD2 | Hydrogen Bond;Electrostatic, Salt Bridge;Attractive Charge | 2.34363 |
| B:LYS126:HZ3 | A:ASP62:OD1 | H-Donor;Positive | H-Acceptor;Negative | B:LYS126:HZ3 - A:ASP62:OD1 | Hydrogen Bond;Electrostatic, Salt Bridge;Attractive Charge | 1.84177 |
| A:LYS164:NZ | B:ASP43:OD1 | Positive | Negative | A:LYS164:NZ - B:ASP43:OD1 | Electrostatic; Attractive Charge | 4.86115 |
| A:LYS164:HZ1 | B:ASP43:O | H-Donor | H-Acceptor | A:LYS164:HZ1 - B:ASP43:O | Conventional Hydrogen Bond | 1.52761 |
| A:LEU166:O1 | B:SER3:O | H-Donor | H-Acceptor | A:LEU166:O1 - B:SER3:O | Conventional Hydrogen Bond | 3.27287 |
| B:MET1:H2 | A:LEU166:O1 | H-Donor | H-Acceptor | B:MET1:H2 - A:LEU166:O1 | Conventional Hydrogen Bond | 1.64651 |
| B:ALA2:H | A:LEU166:O1 | H-Donor | H-Acceptor | B:ALA2:H - A:LEU166:O1 | Conventional Hydrogen Bond | 1.58773 |
| B:SER3:H | A:LEU166:O1 | H-Donor | H-Acceptor | B:SER3:H - A:LEU166:O1 | Conventional Hydrogen Bond | 2.02605 |
| B:ARG45:HH11 | A:PRO165:O | H-Donor | H-Acceptor | B:ARG45:HH11 - A:PRO165:O | Conventional Hydrogen Bond | 2.55151 |
| B:SER119:HG | A:GLN26:OE1 | H-Donor | H-Acceptor | B:SER119:HG - A:GLN26:OE1 | Conventional Hydrogen Bond | 2.7524 |
| A:LYS30:HE1 | B:ASP122:OD2 | H-Donor | H-Acceptor | A:LYS30:HE1 - B:ASP122:OD2 | Carbon Hydrogen Bond | 2.79918 |
| A:LYS30:HE2 | B:ASP122:OD2 | H-Donor | H-Acceptor | A:LYS30:HE2 - B:ASP122:OD2 | Carbon Hydrogen Bond | 2.53537 |
| A:LEU166:HA | B:SER3:O | H-Donor | H-Acceptor | A:LEU166:HA - B:SER3:O | Carbon Hydrogen Bond | 3.07275 |
| B:SER3:HB2 | A:PRO165:O | H-Donor | H-Acceptor | B:SER3:HB2 - A:PRO165:O | Carbon Hydrogen Bond | 2.43197 |
| B:ARG45:HD2 | A:PRO165:O | H-Donor | H-Acceptor | B:ARG45:HD2 - A:PRO165:O | Carbon Hydrogen Bond | 2.56583 |
| B:SER119:HB1 | A:GLN26:OE1 | H-Donor | H-Acceptor | B:SER119:HB1 - A:GLN26:OE1 | Carbon Hydrogen Bond | 3.09079 |
| B:LYS126:HE2 | A:ASP59:OD2 | H-Donor | H-Acceptor | B:LYS126:HE2 - A:ASP59:OD2 | Carbon Hydrogen Bond | 2.6808 |
| A:LYS34 | B:MET1 | Alkyl | Alkyl | A:LYS34 - B:MET1 | Hydrophobic Alkyl | 5.16168 |
| A:PRO67 | B:MET1 | Alkyl | Alkyl | A:PRO67 - B:MET1 | Hydrophobic Alkyl | 4.98526 |
| B:ALA2 | A:PRO165 | Alkyl | Alkyl | B:ALA2 - A:PRO165 | Hydrophobic Alkyl | 4.59526 |
| B:ARG45 | A:LYS164 | Alkyl | Alkyl | B:ARG45 - A:LYS164 | Hydrophobic Alkyl | 5.33126 |

D. **Complex 9**

| **From** | **To** | **From Chemistry** | **To Chemistry** | **Interaction** | **Type** | **Distance** |
| --- | --- | --- | --- | --- | --- | --- |
| B:LYS112:HZ2 | A:ASP145:OD1 | H-Donor;Positive | H-Acceptor;Negative | B:LYS112:HZ2 - A:ASP145:OD1 | Hydrogen Bond;Electrostatic; Salt Bridge;Attractive Charge | 1.77056 |
| B:ARG45:NH1 | A:ASP66:OD1 | Positive | Negative | B:ARG45:NH1 - A:ASP66:OD1 | Electrostatic, Attractive Charge | 4.62484 |
| B:LYS121:NZ | A:GLU162:OE2 | Positive | Negative | B:LYS121:NZ - A:GLU162:OE2 | Electrostatic, Attractive Charge | 5.18603 |
| B:LYS126:NZ | A:GLU65:OE2 | Positive | Negative | B:LYS126:NZ - A:GLU65:OE2 | Electrostatic, Attractive Charge | 4.92253 |
| A:THR148:H | B:MET1:O | H-Donor | H-Acceptor | A:THR148:H - B:MET1:O | Conventional Hydrogen Bond | 2.02793 |
| B:MET1:H3 | A:THR148:OG1 | H-Donor | H-Acceptor | B:MET1:H3 - A:THR148:OG1 | Conventional Hydrogen Bond | 1.96 |
| B:MET1:H3 | A:THR148:O | H-Donor | H-Acceptor | B:MET1:H3 - A:THR148:O | Conventional Hydrogen Bond | 2.41216 |
| B:SER3:HG | A:ARG146:O | H-Donor | H-Acceptor | B:SER3:HG - A:ARG146:O | Conventional Hydrogen Bond | 2.98158 |
| B:ARG45:HH11 | A:THR69:OG1 | H-Donor | H-Acceptor | B:ARG45:HH11 - A:THR69:OG1 | Conventional Hydrogen Bond | 1.68932 |
| A:ARG146:HA | B:MET1:O | H-Donor | H-Acceptor | A:ARG146:HA - B:MET1:O | Carbon Hydrogen Bond | 2.63505 |
| B:ARG45:HD2 | A:ASP66:OD1 | H-Donor | H-Acceptor | B:ARG45:HD2 - A:ASP66:OD1 | Carbon Hydrogen Bond | 2.68457 |
| B:LYS112:HE2 | A:ASP145:OD1 | H-Donor | H-Acceptor | B:LYS112:HE2 - A:ASP145:OD1 | Carbon Hydrogen Bond | 2.95603 |
| B:LYS112:HE2 | A:ASP145:O | H-Donor | H-Acceptor | B:LYS112:HE2 - A:ASP145:O | Carbon Hydrogen Bond | 2.89548 |
| B:MET115:HA | A:GLY163:O | H-Donor | H-Acceptor | B:MET115:HA - A:GLY163:O | Carbon Hydrogen Bond | 2.53176 |
| B:MET1:SD | A:TRP135 | Sulfur | Pi-Orbitals | B:MET1:SD - A:TRP135 | Pi-Sulfur | 5.47455 |
| A:PRO67 | B:MET1 | Alkyl | Alkyl | A:PRO67 - B:MET1 | Hydrophobic, Alkyl | 5.11322 |
| A:ILE143 | B:MET1 | Alkyl | Alkyl | A:ILE143 - B:MET1 | Hydrophobic, Alkyl | 5.40409 |
| A:PRO165 | B:LYS114 | Alkyl | Alkyl | A:PRO165 - B:LYS114 | Hydrophobic, Alkyl | 4.61165 |
| B:ALA2 | A:LEU149 | Alkyl | Alkyl | B:ALA2 - A:LEU149 | Hydrophobic, Alkyl | 5.36983 |
| A:PHE71 | B:MET1 | Pi-Orbitals | Alkyl | A:PHE71 - B:MET1 | Hydrophobic, Pi-Alkyl | 5.32325 |

**Table S4:** The table denotes solvent accessibility of APRs in stable cofilin-2 homodimers (A) Complex 1, (B) Complex 3, (C) Complex 7, and (D) Complex 9. Blue, Cyan, and Green represent % residue solvent accessibility of more than 25%, between 10% to 25%, and less than 10%, respectively.

A. **Complex 1**

| **Residue name** | **Hydrophobicity** | **pKa** | **Secondary structure** | **Residue solvent accessibility** | **Side chain solvent accessibility** | **% residue solvent accessibility** | **% side chain solvent accessibility** |
| --- | --- | --- | --- | --- | --- | --- | --- |
| A:Met1 | 1.9 |  | Coil | 92.707 | 31.061 | 50.948 | 24.075 |
| A:Ala2 | 1.8 |  | Coil | 33.937 | 12.579 | 33.896 | 27.322 |
| A:Val7 | 4.2 |  | Coil | 21.365 | 2.516 | 15.193 | 2.861 |
| A:Ile12 | 4.5 |  | Helix | 53.145 | 49.311 | 33.262 | 43.459 |
| A:Lys13 | -3.9 | 10.4 | Helix | 113.789 | 93.99 | 58.514 | 66.414 |
| A:Val14 | 4.2 |  | Helix | 16.102 | 14.592 | 11.45 | 16.595 |
| A:Phe15 | 2.8 |  | Helix | 8.035 | 7.548 | 4.017 | 5.068 |
| A:Ile29 | 4.5 |  | Helix | 75.692 | 70.445 | 47.373 | 62.084 |
| A:Ala35 | 1.8 |  | Sheet | 0.503 | 0.503 | 0.503 | 1.093 |
| A:Val36 | 4.2 |  | Sheet | 0.503 | 0.503 | 0.358 | 0.572 |
| A:Leu37 | 3.8 |  | Sheet | 1.006 | 1.006 | 0.663 | 1.013 |
| A:Phe38 | 2.8 |  | Sheet | 0.503 | 0.503 | 0.252 | 0.338 |
| A:Cys39 | 2.5 | 9 | Sheet | 12.712 | 12.712 | 9.79 | 16.423 |
| A:Leu40 | 3.8 |  | Sheet | 19.656 | 4.529 | 12.957 | 4.557 |
| A:Ser41 | -0.8 |  | Coil | 17.699 | 15.343 | 15.597 | 25.348 |
| A:Gln46 | -3.5 |  | Coil | 74.039 | 74.039 | 41.524 | 59.063 |
| A:Ile47 | 4.5 |  | Sheet | 0.503 | 0.503 | 0.315 | 0.443 |
| A:Ile48 | 4.5 |  | Sheet | 48.808 | 47.802 | 30.547 | 42.129 |
| A:Val49 | 4.2 |  | Sheet | 33.026 | 7.044 | 23.485 | 8.011 |
| A:Glu50 | -3.5 | 4.3 | Sheet | 61.956 | 60.446 | 35.129 | 48.975 |
| A:Gln54 | -3.5 |  | Sheet | 40.88 | 28.628 | 22.927 | 22.837 |
| A:Ile55 | 4.5 |  | Sheet | 2.013 | 2.013 | 1.26 | 1.774 |
| A:Leu56 | 3.8 |  | Coil | 55.334 | 54.846 | 36.477 | 55.19 |
| A:Val57 | 4.2 |  | Turn | 98.271 | 76.986 | 69.882 | 87.554 |
| A:Gly58 | -0.4 |  | Turn | 61.826 | 25.662 | 86.314 | 106.806 |
| A:Asp59 | -3.5 | 3.9 | Turn | 31.74 | 28.25 | 21.76 | 29.838 |
| A:Val64 | 4.2 |  | Coil | 17.595 | 15.599 | 12.512 | 17.74 |
| A:Tyr68 | -1.3 | 10 | Turn | 54.799 | 54.799 | 24.958 | 32.52 |
| A:Thr69 | -0.7 |  | Helix | 15.247 | 10.064 | 11.499 | 11.663 |
| A:Ser70 | -0.8 |  | Helix | 6.693 | 3.834 | 5.898 | 6.334 |
| A:Phe71 | 2.8 |  | Helix | 4.025 | 3.522 | 2.013 | 2.365 |
| A:Val72 | 4.2 |  | Helix | 48.393 | 41.764 | 34.413 | 47.496 |
| A:Lys73 | -3.9 | 10.4 | Turn | 53.883 | 42.613 | 27.708 | 30.111 |
| A:Leu74 | 3.8 |  | Turn | 5.407 | 3.522 | 3.565 | 3.544 |
| A:Leu75 | 3.8 |  | Coil | 9.992 | 1.51 | 6.587 | 1.519 |
| A:Pro76 | -1.6 |  | Coil | 33.697 | 25.662 | 26.49 | 27.383 |
| A:Leu77 | 3.8 |  | Turn | 124.435 | 100.636 | 82.03 | 101.266 |
| A:Arg81 | -4.5 | 12 | Sheet | 40.524 | 40.524 | 17.666 | 22.968 |
| A:Tyr82 | -1.3 | 10 | Sheet | 1.006 | 0 | 0.458 | 0 |
| A:Ala83 | 1.8 |  | Sheet | 4.529 | 4.529 | 4.523 | 9.836 |
| A:Leu84 | 3.8 |  | Sheet | 1.006 | 1.006 | 0.663 | 1.013 |
| A:Tyr85 | -1.3 | 10 | Sheet | 15.998 | 15.998 | 7.286 | 9.494 |
| A:Asp86 | -3.5 | 3.9 | Sheet | 34.185 | 32.3 | 23.436 | 34.115 |
| A:Ala87 | 1.8 |  | Sheet | 9.968 | 1.51 | 9.956 | 3.279 |
| A:Asp98 | -3.5 | 3.9 | Sheet | 17.627 | 15.615 | 12.085 | 16.492 |
| A:Leu99 | 3.8 |  | Sheet | 10.47 | 7.548 | 6.902 | 7.595 |
| A:Val100 | 4.2 |  | Sheet | 1.51 | 1.51 | 1.073 | 1.717 |
| A:Phe101 | 2.8 |  | Sheet | 1.006 | 1.006 | 0.503 | 0.676 |
| A:Ile102 | 4.5 |  | Sheet | 4.465 | 3.522 | 2.794 | 3.104 |
| A:Phe103 | 2.8 |  | Sheet | 0 | 0 | 0 | 0 |
| A:Trp104 | -0.9 |  | Sheet | 4.529 | 4.529 | 1.87 | 2.367 |
| A:Ala105 | 1.8 |  | Coil | 9.744 | 5.032 | 9.733 | 10.929 |
| A:Pro110 | -1.6 |  | Coil | 49.766 | 39.751 | 39.122 | 42.416 |
| A:Leu111 | 3.8 |  | Helix | 142.701 | 127.807 | 94.071 | 128.608 |
| A:Ser113 | -0.8 |  | Helix | 5.216 | 5.216 | 4.596 | 8.617 |
| A:Lys114 | -3.9 | 10.4 | Helix | 57.455 | 57.455 | 29.545 | 40.598 |
| A:Met115 | 1.9 |  | Helix | 110.011 | 105.091 | 60.458 | 81.455 |
| A:Ile116 | 4.5 |  | Turn | 20.111 | 19.121 | 12.587 | 16.851 |
| A:Tyr117 | -1.3 | 10 | Turn | 0.471 | 0.471 | 0.215 | 0.28 |
| A:Ala118 | 1.8 |  | Turn | 65.757 | 32.707 | 65.679 | 71.038 |
| A:Lys126 | -3.9 | 10.4 | Turn | 126.081 | 108.453 | 64.834 | 76.634 |
| A:Lys127 | -3.9 | 10.4 | Turn | 97.952 | 85.979 | 50.37 | 60.754 |
| A:Phe128 | 2.8 |  | Coil | 0 | 0 | 0 | 0 |
| A:Thr129 | -0.7 |  | Turn | 88.991 | 74.567 | 67.115 | 86.42 |
| A:Gly130 | -0.4 |  | Turn | 75.435 | 29.688 | 105.312 | 123.56 |
| A:Gly155 | -0.4 |  | Coil | 42.02 | 13.083 | 58.663 | 54.45 |
| A:Asn156 | -3.5 |  | Coil | 144.761 | 121.695 | 98.074 | 125.889 |
| A:Val157 | 4.2 |  | Coil | 68.416 | 63.904 | 48.651 | 72.675 |
| A:Val158 | 4.2 |  | Turn | 69.262 | 63.904 | 49.253 | 72.675 |
| A:Val159 | 4.2 |  | Turn | 105.332 | 85.037 | 74.903 | 96.71 |
| A:Ser160 | -0.8 |  | Turn | 23.29 | 3.522 | 20.524 | 5.819 |
| A:Leu161 | 3.8 |  | Turn | 38.242 | 37.738 | 25.209 | 37.975 |
| A:Pro165 | -1.6 |  | Turn | 52.227 | 43.273 | 41.056 | 46.174 |
| A:Leu166 | 3.8 |  | Coil | 47.012 | 12.579 | 30.991 | 12.658 |
| B:Met1 | 1.9 |  | Coil | 17.092 | 16.605 | 9.393 | 12.87 |
| B:Ala2 | 1.8 |  | Coil | 7.532 | 4.529 | 7.523 | 9.836 |
| B:Val7 | 4.2 |  | Coil | 8.985 | 0 | 6.39 | 0 |
| B:Ile12 | 4.5 |  | Helix | 32.707 | 32.707 | 20.47 | 28.825 |
| B:Lys13 | -3.9 | 10.4 | Helix | 113.845 | 107.559 | 58.542 | 76.003 |
| B:Val14 | 4.2 |  | Helix | 24.528 | 22.14 | 17.442 | 25.179 |
| B:Phe15 | 2.8 |  | Helix | 1.006 | 0 | 0.503 | 0 |
| B:Ile29 | 4.5 |  | Helix | 95.652 | 80.005 | 59.865 | 70.51 |
| B:Ala35 | 1.8 |  | Sheet | 0 | 0 | 0 | 0 |
| B:Val36 | 4.2 |  | Sheet | 1.51 | 1.51 | 1.073 | 1.717 |
| B:Leu37 | 3.8 |  | Sheet | 7.044 | 7.044 | 4.644 | 7.089 |
| B:Phe38 | 2.8 |  | Sheet | 1.006 | 1.006 | 0.503 | 0.676 |
| B:Cys39 | 2.5 | 9 | Sheet | 12.846 | 12.846 | 9.893 | 16.596 |
| B:Leu40 | 3.8 |  | Sheet | 39.903 | 19.624 | 26.305 | 19.747 |
| B:Ser41 | -0.8 |  | Coil | 25.151 | 11.573 | 22.164 | 19.12 |
| B:Gln46 | -3.5 |  | Coil | 85.081 | 85.081 | 47.717 | 67.87 |
| B:Ile47 | 4.5 |  | Sheet | 2.516 | 2.516 | 1.575 | 2.217 |
| B:Ile48 | 4.5 |  | Sheet | 63.904 | 62.897 | 39.995 | 55.432 |
| B:Val49 | 4.2 |  | Coil | 34.927 | 6.541 | 24.837 | 7.439 |
| B:Glu50 | -3.5 | 4.3 | Coil | 110.709 | 107.25 | 62.771 | 86.897 |
| B:Gln54 | -3.5 |  | Sheet | 50.919 | 25.928 | 28.558 | 20.683 |
| B:Ile55 | 4.5 |  | Sheet | 10.064 | 5.032 | 6.298 | 4.435 |
| B:Leu56 | 3.8 |  | Coil | 53.192 | 48.808 | 35.065 | 49.114 |
| B:Val57 | 4.2 |  | Turn | 54.463 | 48.808 | 38.729 | 55.508 |
| B:Gly58 | -0.4 |  | Turn | 42.667 | 16.102 | 59.566 | 67.016 |
| B:Asp59 | -3.5 | 3.9 | Turn | 91.043 | 82.634 | 62.417 | 87.279 |
| B:Val64 | 4.2 |  | Coil | 42.738 | 34.719 | 30.392 | 39.485 |
| B:Tyr68 | -1.3 | 10 | Helix | 119.206 | 119.206 | 54.292 | 70.743 |
| B:Thr69 | -0.7 |  | Helix | 56.707 | 56.22 | 42.767 | 65.158 |
| B:Ser70 | -0.8 |  | Helix | 23.681 | 12.036 | 20.869 | 19.885 |
| B:Phe71 | 2.8 |  | Helix | 1.51 | 1.51 | 0.755 | 1.014 |
| B:Val72 | 4.2 |  | Helix | 43.729 | 27.675 | 31.096 | 31.474 |
| B:Lys73 | -3.9 | 10.4 | Helix | 161.157 | 129.703 | 82.871 | 91.65 |
| B:Leu74 | 3.8 |  | Helix | 10.551 | 7.044 | 6.955 | 7.089 |
| B:Leu75 | 3.8 |  | Coil | 0 | 0 | 0 | 0 |
| B:Pro76 | -1.6 |  | Coil | 34.216 | 33.713 | 26.898 | 35.973 |
| B:Leu77 | 3.8 |  | Turn | 97.201 | 74.47 | 64.076 | 74.937 |
| B:Arg81 | -4.5 | 12 | Sheet | 56.046 | 56.046 | 24.433 | 31.765 |
| B:Tyr82 | -1.3 | 10 | Sheet | 4.529 | 4.529 | 2.063 | 2.688 |
| B:Ala83 | 1.8 |  | Sheet | 0 | 0 | 0 | 0 |
| B:Leu84 | 3.8 |  | Sheet | 0.503 | 0.503 | 0.332 | 0.506 |
| B:Tyr85 | -1.3 | 10 | Sheet | 2.42 | 2.42 | 1.102 | 1.436 |
| B:Asp86 | -3.5 | 3.9 | Sheet | 23.091 | 23.091 | 15.83 | 24.388 |
| B:Ala87 | 1.8 |  | Sheet | 0 | 0 | 0 | 0 |
| B:Asp98 | -3.5 | 3.9 | Sheet | 12.995 | 8.482 | 8.909 | 8.959 |
| B:Leu99 | 3.8 |  | Sheet | 21.053 | 18.618 | 13.878 | 18.734 |
| B:Val100 | 4.2 |  | Sheet | 0 | 0 | 0 | 0 |
| B:Phe101 | 2.8 |  | Sheet | 2.013 | 2.013 | 1.006 | 1.351 |
| B:Ile102 | 4.5 |  | Sheet | 0 | 0 | 0 | 0 |
| B:Phe103 | 2.8 |  | Sheet | 31.7 | 31.7 | 15.85 | 21.284 |
| B:Trp104 | -0.9 |  | Sheet | 0 | 0 | 0 | 0 |
| B:Ala105 | 1.8 |  | Coil | 19.329 | 7.548 | 19.306 | 16.393 |
| B:Pro110 | -1.6 |  | Coil | 54.343 | 48.808 | 42.72 | 52.081 |
| B:Leu111 | 3.8 |  | Helix | 128.932 | 120.763 | 84.994 | 121.519 |
| B:Ser113 | -0.8 |  | Helix | 20.838 | 19.424 | 18.363 | 32.091 |
| B:Lys114 | -3.9 | 10.4 | Helix | 58.671 | 58.671 | 30.17 | 41.457 |
| B:Met115 | 1.9 |  | Helix | 77.358 | 72.926 | 42.513 | 56.524 |
| B:Ile116 | 4.5 |  | Turn | 6.725 | 2.013 | 4.209 | 1.774 |
| B:Tyr117 | -1.3 | 10 | Turn | 13.266 | 10.846 | 6.042 | 6.437 |
| B:Ala118 | 1.8 |  | Helix | 39.424 | 26.668 | 39.377 | 57.924 |
| B:Lys126 | -3.9 | 10.4 | Helix | 170.353 | 148.325 | 87.6 | 104.809 |
| B:Lys127 | -3.9 | 10.4 | Helix | 64.517 | 51.57 | 33.176 | 36.44 |
| B:Phe128 | 2.8 |  | Coil | 0 | 0 | 0 | 0 |
| B:Thr129 | -0.7 |  | Turn | 69.671 | 63.696 | 52.544 | 73.822 |
| B:Gly130 | -0.4 |  | Turn | 44.165 | 20.63 | 61.657 | 85.864 |
| B:Gly155 | -0.4 |  | Coil | 33.338 | 3.019 | 46.542 | 12.565 |
| B:Asn156 | -3.5 |  | Coil | 122.517 | 97.311 | 83.005 | 100.664 |
| B:Val157 | 4.2 |  | Coil | 60.381 | 60.381 | 42.938 | 68.67 |
| B:Val158 | 4.2 |  | Sheet | 13.706 | 7.548 | 9.746 | 8.584 |
| B:Val159 | 4.2 |  | Sheet | 87.306 | 56.859 | 62.084 | 64.664 |
| B:Ser160 | -0.8 |  | Sheet | 21.82 | 19.808 | 19.229 | 32.724 |
| B:Leu161 | 3.8 |  | Sheet | 14.552 | 5.535 | 9.593 | 5.57 |
| B:Pro165 | -1.6 |  | Coil | 76.675 | 49.311 | 60.275 | 52.617 |
| B:Leu166 | 3.8 |  | Coil | 223.771 | 124.788 | 147.513 | 125.57 |

B. **Complex 3**

| **Residue name** | **Hydrophobicity** | **pKa** | **Secondary structure** | **Residue solvent accessibility** | **Side chain solvent accessibility** | **% residue solvent accessibility** | **% side chain solvent accessibility** |
| --- | --- | --- | --- | --- | --- | --- | --- |
| A:Met1 | 1.9 |  | Coil | 164.871 | 119.116 | 90.607 | 92.326 |
| A:Ala2 | 1.8 |  | Coil | 14.177 | 1.006 | 14.16 | 2.186 |
| A:Val7 | 4.2 |  | Coil | 7.651 | 0 | 5.44 | 0 |
| A:Ile12 | 4.5 |  | Helix | 58.872 | 58.872 | 36.846 | 51.885 |
| A:Lys13 | -3.9 | 10.4 | Helix | 123.171 | 110.695 | 63.338 | 78.219 |
| A:Val14 | 4.2 |  | Turn | 37.531 | 30.694 | 26.688 | 34.907 |
| A:Phe15 | 2.8 |  | Turn | 8.554 | 8.554 | 4.277 | 5.743 |
| A:Ile29 | 4.5 |  | Helix | 44.248 | 42.77 | 27.693 | 37.694 |
| A:Ala35 | 1.8 |  | Sheet | 1.006 | 1.006 | 1.005 | 2.186 |
| A:Val36 | 4.2 |  | Sheet | 3.019 | 3.019 | 2.147 | 3.433 |
| A:Leu37 | 3.8 |  | Sheet | 16.605 | 16.605 | 10.946 | 16.709 |
| A:Phe38 | 2.8 |  | Sheet | 2.356 | 0 | 1.178 | 0 |
| A:Cys39 | 2.5 | 9 | Sheet | 19.119 | 19.119 | 14.724 | 24.699 |
| A:Leu40 | 3.8 |  | Sheet | 11.374 | 1.006 | 7.498 | 1.013 |
| A:Ser41 | -0.8 |  | Coil | 19.177 | 13.802 | 16.899 | 22.801 |
| A:Gln46 | -3.5 |  | Coil | 55.64 | 55.64 | 31.205 | 44.385 |
| A:Ile47 | 4.5 |  | Sheet | 0 | 0 | 0 | 0 |
| A:Ile48 | 4.5 |  | Sheet | 49.815 | 49.815 | 31.177 | 43.902 |
| A:Val49 | 4.2 |  | Sheet | 21.437 | 11.07 | 15.244 | 12.589 |
| A:Glu50 | -3.5 | 4.3 | Coil | 53.729 | 44.304 | 30.464 | 35.897 |
| A:Gln54 | -3.5 |  | Coil | 34.118 | 16.947 | 19.135 | 13.519 |
| A:Ile55 | 4.5 |  | Sheet | 22.643 | 17.108 | 14.172 | 15.078 |
| A:Leu56 | 3.8 |  | Sheet | 78.055 | 61.388 | 51.455 | 61.772 |
| A:Val57 | 4.2 |  | Coil | 47.451 | 39.751 | 33.743 | 45.207 |
| A:Gly58 | -0.4 |  | Coil | 27.33 | 11.07 | 38.154 | 46.073 |
| A:Asp59 | -3.5 | 3.9 | Coil | 79.223 | 74.807 | 54.313 | 79.012 |
| A:Val64 | 4.2 |  | Coil | 27.172 | 27.172 | 19.322 | 30.901 |
| A:Tyr68 | -1.3 | 10 | Turn | 21.102 | 21.102 | 9.611 | 12.523 |
| A:Thr69 | -0.7 |  | Turn | 49.176 | 46.348 | 37.087 | 53.716 |
| A:Ser70 | -0.8 |  | Helix | 27.396 | 26.924 | 24.142 | 44.481 |
| A:Phe71 | 2.8 |  | Helix | 3.019 | 3.019 | 1.51 | 2.027 |
| A:Val72 | 4.2 |  | Helix | 24.672 | 10.064 | 17.544 | 11.445 |
| A:Lys73 | -3.9 | 10.4 | Helix | 143.454 | 102.041 | 73.768 | 72.103 |
| A:Leu74 | 3.8 |  | Turn | 82.785 | 60.885 | 54.573 | 61.266 |
| A:Leu75 | 3.8 |  | Coil | 12.412 | 1.006 | 8.182 | 1.013 |
| A:Pro76 | -1.6 |  | Coil | 32.171 | 24.153 | 25.29 | 25.772 |
| A:Leu77 | 3.8 |  | Coil | 49.526 | 42.77 | 32.648 | 43.038 |
| A:Arg81 | -4.5 | 12 | Sheet | 49.372 | 48.901 | 21.524 | 27.716 |
| A:Tyr82 | -1.3 | 10 | Sheet | 1.478 | 1.006 | 0.673 | 0.597 |
| A:Ala83 | 1.8 |  | Sheet | 1.493 | 1.006 | 1.492 | 2.186 |
| A:Leu84 | 3.8 |  | Sheet | 0 | 0 | 0 | 0 |
| A:Tyr85 | -1.3 | 10 | Sheet | 6.382 | 6.382 | 2.906 | 3.787 |
| A:Asp86 | -3.5 | 3.9 | Sheet | 10.966 | 10.966 | 7.518 | 11.583 |
| A:Ala87 | 1.8 |  | Sheet | 2.484 | 2.013 | 2.481 | 4.372 |
| A:Asp98 | -3.5 | 3.9 | Sheet | 75.581 | 69.088 | 51.816 | 72.971 |
| A:Leu99 | 3.8 |  | Sheet | 21.268 | 13.586 | 14.02 | 13.671 |
| A:Val100 | 4.2 |  | Sheet | 16.102 | 12.579 | 11.45 | 14.306 |
| A:Phe101 | 2.8 |  | Sheet | 6.333 | 1.006 | 3.166 | 0.676 |
| A:Ile102 | 4.5 |  | Sheet | 2.516 | 2.516 | 1.575 | 2.217 |
| A:Phe103 | 2.8 |  | Sheet | 11.573 | 11.573 | 5.787 | 7.77 |
| A:Trp104 | -0.9 |  | Sheet | 0 | 0 | 0 | 0 |
| A:Ala105 | 1.8 |  | Coil | 2.452 | 1.51 | 2.449 | 3.279 |
| A:Pro110 | -1.6 |  | Coil | 31.7 | 31.7 | 24.92 | 33.826 |
| A:Leu111 | 3.8 |  | Helix | 86.107 | 69.439 | 56.763 | 69.873 |
| A:Ser113 | -0.8 |  | Helix | 9.912 | 9.425 | 8.735 | 15.57 |
| A:Lys114 | -3.9 | 10.4 | Helix | 74.906 | 72.015 | 38.519 | 50.887 |
| A:Met115 | 1.9 |  | Helix | 82.188 | 78.73 | 45.167 | 61.023 |
| A:Ile116 | 4.5 |  | Helix | 9.56 | 9.56 | 5.984 | 8.426 |
| A:Tyr117 | -1.3 | 10 | Helix | 5.942 | 5.471 | 2.706 | 3.247 |
| A:Ala118 | 1.8 |  | Helix | 57.275 | 33.713 | 57.207 | 73.224 |
| A:Lys126 | -3.9 | 10.4 | Turn | 142.63 | 115.546 | 73.344 | 81.646 |
| A:Lys127 | -3.9 | 10.4 | Turn | 58.492 | 52.741 | 30.078 | 37.268 |
| A:Phe128 | 2.8 |  | Turn | 0 | 0 | 0 | 0 |
| A:Thr129 | -0.7 |  | Turn | 85.437 | 65.39 | 64.434 | 75.784 |
| A:Gly130 | -0.4 |  | Turn | 76.641 | 26.165 | 106.996 | 108.901 |
| A:Gly155 | -0.4 |  | Coil | 45.77 | 23.649 | 63.898 | 98.429 |
| A:Asn156 | -3.5 |  | Coil | 121.286 | 84.7 | 82.171 | 87.618 |
| A:Val157 | 4.2 |  | Coil | 83.903 | 80.005 | 59.664 | 90.987 |
| A:Val158 | 4.2 |  | Sheet | 46.029 | 34.719 | 32.732 | 39.485 |
| A:Val159 | 4.2 |  | Sheet | 65.781 | 55.853 | 46.777 | 63.519 |
| A:Ser160 | -0.8 |  | Sheet | 19.297 | 19.297 | 17.005 | 31.88 |
| A:Leu161 | 3.8 |  | Sheet | 31.852 | 26.668 | 20.997 | 26.835 |
| A:Pro165 | -1.6 |  | Coil | 69.407 | 68.935 | 54.561 | 73.557 |
| A:Leu166 | 3.8 |  | Coil | 149.485 | 67.426 | 98.543 | 67.848 |
| B:Met1 | 1.9 |  | Coil | 184.26 | 96.44 | 101.262 | 74.75 |
| B:Ala2 | 1.8 |  | Coil | 71.082 | 51.324 | 70.997 | 111.475 |
| B:Val7 | 4.2 |  | Sheet | 14.672 | 0.503 | 10.433 | 0.572 |
| B:Ile12 | 4.5 |  | Helix | 47.738 | 46.796 | 29.878 | 41.242 |
| B:Lys13 | -3.9 | 10.4 | Helix | 116.497 | 104.508 | 59.906 | 73.847 |
| B:Val14 | 4.2 |  | Helix | 25.063 | 20.63 | 17.822 | 23.462 |
| B:Phe15 | 2.8 |  | Helix | 8.554 | 8.554 | 4.277 | 5.743 |
| B:Ile29 | 4.5 |  | Turn | 49.152 | 46.796 | 30.762 | 41.242 |
| B:Ala35 | 1.8 |  | Sheet | 0 | 0 | 0 | 0 |
| B:Val36 | 4.2 |  | Sheet | 1.51 | 1.51 | 1.073 | 1.717 |
| B:Leu37 | 3.8 |  | Sheet | 1.006 | 1.006 | 0.663 | 1.013 |
| B:Phe38 | 2.8 |  | Sheet | 0.471 | 0 | 0.236 | 0 |
| B:Cys39 | 2.5 | 9 | Sheet | 31.899 | 30.39 | 24.566 | 39.26 |
| B:Leu40 | 3.8 |  | Sheet | 42.338 | 18.618 | 27.91 | 18.734 |
| B:Ser41 | -0.8 |  | Coil | 26.125 | 18.49 | 23.023 | 30.547 |
| B:Gln46 | -3.5 |  | Sheet | 80.187 | 79.7 | 44.972 | 63.578 |
| B:Ile47 | 4.5 |  | Sheet | 0 | 0 | 0 | 0 |
| B:Ile48 | 4.5 |  | Sheet | 55.853 | 54.846 | 34.956 | 48.337 |
| B:Val49 | 4.2 |  | Sheet | 20.398 | 8.051 | 14.505 | 9.156 |
| B:Glu50 | -3.5 | 4.3 | Sheet | 39.113 | 39.113 | 22.177 | 31.69 |
| B:Gln54 | -3.5 |  | Sheet | 43.206 | 30.954 | 24.232 | 24.692 |
| B:Ile55 | 4.5 |  | Sheet | 22.643 | 18.618 | 14.172 | 16.408 |
| B:Leu56 | 3.8 |  | Sheet | 55.788 | 53.84 | 36.777 | 54.177 |
| B:Val57 | 4.2 |  | Turn | 45.558 | 34.719 | 32.397 | 39.485 |
| B:Gly58 | -0.4 |  | Turn | 32.771 | 16.605 | 45.75 | 69.11 |
| B:Asp59 | -3.5 | 3.9 | Turn | 86.588 | 85.142 | 59.362 | 89.928 |
| B:Val64 | 4.2 |  | Coil | 27.172 | 27.172 | 19.322 | 30.901 |
| B:Tyr68 | -1.3 | 10 | Helix | 52.435 | 51.46 | 23.881 | 30.539 |
| B:Thr69 | -0.7 |  | Helix | 60.214 | 55.501 | 45.412 | 64.324 |
| B:Ser70 | -0.8 |  | Helix | 0.942 | 0.471 | 0.831 | 0.779 |
| B:Phe71 | 2.8 |  | Helix | 8.019 | 5.535 | 4.009 | 3.716 |
| B:Val72 | 4.2 |  | Turn | 34.767 | 20.63 | 24.723 | 23.462 |
| B:Lys73 | -3.9 | 10.4 | Turn | 82.523 | 68.666 | 42.436 | 48.52 |
| B:Leu74 | 3.8 |  | Turn | 1.51 | 1.51 | 0.995 | 1.519 |
| B:Leu75 | 3.8 |  | Coil | 3.363 | 1.006 | 2.217 | 1.013 |
| B:Pro76 | -1.6 |  | Coil | 52.722 | 27.172 | 41.445 | 28.993 |
| B:Leu77 | 3.8 |  | Coil | 0 | 0 | 0 | 0 |
| B:Arg81 | -4.5 | 12 | Sheet | 30.783 | 30.312 | 13.42 | 17.18 |
| B:Tyr82 | -1.3 | 10 | Sheet | 2.923 | 2.923 | 1.331 | 1.735 |
| B:Ala83 | 1.8 |  | Sheet | 0 | 0 | 0 | 0 |
| B:Leu84 | 3.8 |  | Sheet | 0 | 0 | 0 | 0 |
| B:Tyr85 | -1.3 | 10 | Sheet | 18.514 | 18.514 | 8.432 | 10.987 |
| B:Asp86 | -3.5 | 3.9 | Sheet | 16.342 | 14.928 | 11.203 | 15.767 |
| B:Ala87 | 1.8 |  | Sheet | 5.535 | 1.006 | 5.528 | 2.186 |
| B:Asp98 | -3.5 | 3.9 | Sheet | 68.545 | 63.026 | 46.992 | 66.568 |
| B:Leu99 | 3.8 |  | Sheet | 12.548 | 12.076 | 8.272 | 12.152 |
| B:Val100 | 4.2 |  | Sheet | 0.503 | 0.503 | 0.358 | 0.572 |
| B:Phe101 | 2.8 |  | Sheet | 12.515 | 10.567 | 6.258 | 7.095 |
| B:Ile102 | 4.5 |  | Sheet | 0.503 | 0.503 | 0.315 | 0.443 |
| B:Phe103 | 2.8 |  | Sheet | 4.529 | 3.019 | 2.264 | 2.027 |
| B:Trp104 | -0.9 |  | Sheet | 5.687 | 0 | 2.349 | 0 |
| B:Ala105 | 1.8 |  | Coil | 22.611 | 22.14 | 22.584 | 48.087 |
| B:Pro110 | -1.6 |  | Coil | 57.866 | 52.834 | 45.489 | 56.376 |
| B:Leu111 | 3.8 |  | Turn | 136.887 | 110.699 | 90.238 | 111.392 |
| B:Ser113 | -0.8 |  | Coil | 40.231 | 36.932 | 35.453 | 61.015 |
| B:Lys114 | -3.9 | 10.4 | Coil | 28.27 | 22.615 | 14.537 | 15.98 |
| B:Met115 | 1.9 |  | Coil | 92.182 | 90.234 | 50.66 | 69.94 |
| B:Ile116 | 4.5 |  | Turn | 32.06 | 28.681 | 20.065 | 25.277 |
| B:Tyr117 | -1.3 | 10 | Turn | 13.171 | 13.171 | 5.998 | 7.816 |
| B:Ala118 | 1.8 |  | Turn | 30.382 | 17.108 | 30.346 | 37.158 |
| B:Lys126 | -3.9 | 10.4 | Helix | 137.077 | 115.304 | 70.489 | 81.476 |
| B:Lys127 | -3.9 | 10.4 | Helix | 124.109 | 102.568 | 63.82 | 72.476 |
| B:Phe128 | 2.8 |  | Turn | 0 | 0 | 0 | 0 |
| B:Thr129 | -0.7 |  | Turn | 109.31 | 59.447 | 82.439 | 68.897 |
| B:Gly130 | -0.4 |  | Coil | 51.261 | 18.114 | 71.564 | 75.393 |
| B:Gly155 | -0.4 |  | Coil | 6.006 | 5.535 | 8.385 | 23.037 |
| B:Asn156 | -3.5 |  | Coil | 129.531 | 96.745 | 87.757 | 100.079 |
| B:Val157 | 4.2 |  | Coil | 119.165 | 91.075 | 84.74 | 103.577 |
| B:Val158 | 4.2 |  | Coil | 10.623 | 4.025 | 7.554 | 4.578 |
| B:Val159 | 4.2 |  | Coil | 75.573 | 55.853 | 53.74 | 63.519 |
| B:Ser160 | -0.8 |  | Sheet | 20.622 | 20.622 | 18.173 | 34.07 |
| B:Leu161 | 3.8 |  | Sheet | 5.535 | 5.032 | 3.649 | 5.063 |
| B:Pro165 | -1.6 |  | Coil | 98.847 | 72.458 | 77.705 | 77.315 |
| B:Leu166 | 3.8 |  | Coil | 161.122 | 93.088 | 106.214 | 93.671 |

C. **Complex 7**

| **Residue name** | **Hydrophobicity** | **pKa** | **Secondary structure** | **Residue solvent accessibility** | **Side chain solvent accessibility** | **% residue solvent accessibility** | **% side chain solvent accessibility** |
| --- | --- | --- | --- | --- | --- | --- | --- |
| A:Met1 | 1.9 |  | Coil | 112.251 | 56.856 | 61.689 | 44.069 |
| A:Ala2 | 1.8 |  | Coil | 40.404 | 30.694 | 40.356 | 66.667 |
| A:Val7 | 4.2 |  | Coil | 1.885 | 0 | 1.34 | 0 |
| A:Ile12 | 4.5 |  | Helix | 29.184 | 29.184 | 18.265 | 25.721 |
| A:Lys13 | -3.9 | 10.4 | Turn | 115.506 | 104.005 | 59.397 | 73.491 |
| A:Val14 | 4.2 |  | Turn | 41.732 | 38.745 | 29.676 | 44.063 |
| A:Phe15 | 2.8 |  | Turn | 3.426 | 0 | 1.713 | 0 |
| A:Ile29 | 4.5 |  | Helix | 29.496 | 26.165 | 18.46 | 23.06 |
| A:Ala35 | 1.8 |  | Sheet | 0 | 0 | 0 | 0 |
| A:Val36 | 4.2 |  | Sheet | 3.019 | 3.019 | 2.147 | 3.433 |
| A:Leu37 | 3.8 |  | Sheet | 1.51 | 1.51 | 0.995 | 1.519 |
| A:Phe38 | 2.8 |  | Sheet | 1.51 | 1.51 | 0.755 | 1.014 |
| A:Cys39 | 2.5 | 9 | Sheet | 41.729 | 38.742 | 32.136 | 50.049 |
| A:Leu40 | 3.8 |  | Sheet | 61.651 | 25.662 | 40.641 | 25.823 |
| A:Ser41 | -0.8 |  | Sheet | 37.938 | 30.99 | 33.433 | 51.197 |
| A:Gln46 | -3.5 |  | Sheet | 73.334 | 73.334 | 41.129 | 58.5 |
| A:Ile47 | 4.5 |  | Sheet | 0 | 0 | 0 | 0 |
| A:Ile48 | 4.5 |  | Sheet | 47.802 | 47.802 | 29.918 | 42.129 |
| A:Val49 | 4.2 |  | Sheet | 33.825 | 20.63 | 24.053 | 23.462 |
| A:Glu50 | -3.5 | 4.3 | Sheet | 33.019 | 28.746 | 18.721 | 23.29 |
| A:Gln54 | -3.5 |  | Sheet | 69.412 | 47.735 | 38.929 | 38.079 |
| A:Ile55 | 4.5 |  | Sheet | 1.51 | 1.51 | 0.945 | 1.33 |
| A:Leu56 | 3.8 |  | Coil | 30.694 | 28.681 | 20.234 | 28.861 |
| A:Val57 | 4.2 |  | Coil | 46.899 | 24.153 | 33.351 | 27.468 |
| A:Gly58 | -0.4 |  | Coil | 25.247 | 17.108 | 35.246 | 71.204 |
| A:Asp59 | -3.5 | 3.9 | Coil | 76.648 | 60.997 | 52.548 | 64.425 |
| A:Val64 | 4.2 |  | Coil | 7.044 | 7.044 | 5.009 | 8.011 |
| A:Tyr68 | -1.3 | 10 | Turn | 34.704 | 34.704 | 15.806 | 20.595 |
| A:Thr69 | -0.7 |  | Helix | 69.95 | 57.666 | 52.755 | 66.833 |
| A:Ser70 | -0.8 |  | Helix | 14.743 | 4.529 | 12.993 | 7.482 |
| A:Phe71 | 2.8 |  | Helix | 4.529 | 4.529 | 2.264 | 3.041 |
| A:Val72 | 4.2 |  | Helix | 39.36 | 26.165 | 27.989 | 29.757 |
| A:Lys73 | -3.9 | 10.4 | Turn | 150.153 | 113.356 | 77.213 | 80.099 |
| A:Leu74 | 3.8 |  | Turn | 14.776 | 7.044 | 9.741 | 7.089 |
| A:Leu75 | 3.8 |  | Coil | 8.546 | 1.006 | 5.634 | 1.013 |
| A:Pro76 | -1.6 |  | Coil | 37.706 | 29.184 | 29.641 | 31.141 |
| A:Leu77 | 3.8 |  | Turn | 99.588 | 73.464 | 65.65 | 73.924 |
| A:Arg81 | -4.5 | 12 | Sheet | 35.637 | 35.637 | 15.536 | 20.198 |
| A:Tyr82 | -1.3 | 10 | Sheet | 0.471 | 0.471 | 0.215 | 0.28 |
| A:Ala83 | 1.8 |  | Sheet | 1.51 | 1.51 | 1.508 | 3.279 |
| A:Leu84 | 3.8 |  | Sheet | 2.452 | 1.51 | 1.616 | 1.519 |
| A:Tyr85 | -1.3 | 10 | Sheet | 5.687 | 5.687 | 2.59 | 3.375 |
| A:Asp86 | -3.5 | 3.9 | Sheet | 6.757 | 6.757 | 4.632 | 7.137 |
| A:Ala87 | 1.8 |  | Sheet | 1.006 | 1.006 | 1.005 | 2.186 |
| A:Asp98 | -3.5 | 3.9 | Sheet | 24.984 | 17.092 | 17.128 | 18.053 |
| A:Leu99 | 3.8 |  | Sheet | 5 | 4.025 | 3.296 | 4.051 |
| A:Val100 | 4.2 |  | Sheet | 0 | 0 | 0 | 0 |
| A:Phe101 | 2.8 |  | Sheet | 0.503 | 0.503 | 0.252 | 0.338 |
| A:Ile102 | 4.5 |  | Sheet | 1.006 | 1.006 | 0.63 | 0.887 |
| A:Phe103 | 2.8 |  | Sheet | 24.656 | 24.656 | 12.328 | 16.554 |
| A:Trp104 | -0.9 |  | Sheet | 0 | 0 | 0 | 0 |
| A:Ala105 | 1.8 |  | Coil | 16.222 | 10.567 | 16.202 | 22.951 |
| A:Pro110 | -1.6 |  | Coil | 55.301 | 44.783 | 43.473 | 47.785 |
| A:Leu111 | 3.8 |  | Helix | 143.021 | 133.845 | 94.282 | 134.684 |
| A:Ser113 | -0.8 |  | Helix | 23.13 | 11.941 | 20.384 | 19.727 |
| A:Lys114 | -3.9 | 10.4 | Helix | 52.826 | 52.355 | 27.165 | 36.995 |
| A:Met115 | 1.9 |  | Turn | 119.116 | 116.6 | 65.462 | 90.376 |
| A:Ile116 | 4.5 |  | Turn | 34.719 | 34.719 | 21.73 | 30.599 |
| A:Tyr117 | -1.3 | 10 | Turn | 4.84 | 4.84 | 2.204 | 2.872 |
| A:Ala118 | 1.8 |  | Turn | 45.774 | 30.191 | 45.719 | 65.574 |
| A:Lys126 | -3.9 | 10.4 | Turn | 115.005 | 85.029 | 59.139 | 60.083 |
| A:Lys127 | -3.9 | 10.4 | Turn | 69.534 | 64.318 | 35.756 | 45.448 |
| A:Phe128 | 2.8 |  | Coil | 1.51 | 1.51 | 0.755 | 1.014 |
| A:Thr129 | -0.7 |  | Turn | 71.587 | 63.728 | 53.989 | 73.859 |
| A:Gly130 | -0.4 |  | Turn | 39.908 | 20.127 | 55.714 | 83.77 |
| A:Gly155 | -0.4 |  | Coil | 21.621 | 21.133 | 30.184 | 87.958 |
| A:Asn156 | -3.5 |  | Turn | 155.317 | 122.628 | 105.226 | 126.853 |
| A:Val157 | 4.2 |  | Turn | 65.501 | 52.331 | 46.578 | 59.514 |
| A:Val158 | 4.2 |  | Coil | 12.883 | 2.516 | 9.161 | 2.861 |
| A:Val159 | 4.2 |  | Coil | 100.564 | 57.362 | 71.512 | 65.236 |
| A:Ser160 | -0.8 |  | Coil | 42.786 | 37.251 | 37.705 | 61.542 |
| A:Leu161 | 3.8 |  | Coil | 70.325 | 43.777 | 46.359 | 44.051 |
| A:Pro165 | -1.6 |  | Coil | 42.267 | 42.267 | 33.227 | 45.101 |
| A:Leu166 | 3.8 |  | Coil | 98.192 | 76.986 | 64.73 | 77.468 |
| B:Met1 | 1.9 |  | Coil | 40.237 | 25.359 | 22.112 | 19.656 |
| B:Ala2 | 1.8 |  | Coil | 62.77 | 29.688 | 62.695 | 64.481 |
| B:Val7 | 4.2 |  | Sheet | 21.493 | 3.522 | 15.284 | 4.006 |
| B:Ile12 | 4.5 |  | Helix | 57.802 | 56.859 | 36.176 | 50.111 |
| B:Lys13 | -3.9 | 10.4 | Helix | 106.234 | 95.077 | 54.629 | 67.182 |
| B:Val14 | 4.2 |  | Helix | 24.624 | 23.649 | 17.51 | 26.896 |
| B:Phe15 | 2.8 |  | Helix | 12.563 | 11.573 | 6.282 | 7.77 |
| B:Ile29 | 4.5 |  | Helix | 54.846 | 54.846 | 34.327 | 48.337 |
| B:Ala35 | 1.8 |  | Sheet | 3.522 | 3.522 | 3.518 | 7.65 |
| B:Val36 | 4.2 |  | Sheet | 3.522 | 3.019 | 2.505 | 3.433 |
| B:Leu37 | 3.8 |  | Sheet | 2.516 | 2.516 | 1.659 | 2.532 |
| B:Phe38 | 2.8 |  | Sheet | 10.064 | 10.064 | 5.032 | 6.757 |
| B:Cys39 | 2.5 | 9 | Sheet | 18.652 | 18.18 | 14.364 | 23.487 |
| B:Leu40 | 3.8 |  | Sheet | 35.135 | 11.573 | 23.162 | 11.646 |
| B:Ser41 | -0.8 |  | Coil | 37.946 | 26.253 | 33.44 | 43.372 |
| B:Gln46 | -3.5 |  | Sheet | 52.718 | 50.77 | 29.566 | 40.5 |
| B:Ile47 | 4.5 |  | Sheet | 0.503 | 0.503 | 0.315 | 0.443 |
| B:Ile48 | 4.5 |  | Sheet | 55.35 | 54.846 | 34.641 | 48.337 |
| B:Val49 | 4.2 |  | Sheet | 29.416 | 9.56 | 20.918 | 10.873 |
| B:Glu50 | -3.5 | 4.3 | Coil | 38.674 | 36.757 | 21.928 | 29.781 |
| B:Gln54 | -3.5 |  | Coil | 40.076 | 23.111 | 22.476 | 18.436 |
| B:Ile55 | 4.5 |  | Sheet | 4.025 | 4.025 | 2.519 | 3.548 |
| B:Leu56 | 3.8 |  | Sheet | 60.381 | 60.381 | 39.804 | 60.759 |
| B:Val57 | 4.2 |  | Turn | 54.927 | 40.757 | 39.059 | 46.352 |
| B:Gly58 | -0.4 |  | Turn | 24.48 | 6.541 | 34.176 | 27.225 |
| B:Asp59 | -3.5 | 3.9 | Coil | 30.662 | 30.662 | 21.021 | 32.386 |
| B:Val64 | 4.2 |  | Coil | 3.426 | 0.503 | 2.437 | 0.572 |
| B:Tyr68 | -1.3 | 10 | Helix | 69.998 | 69.998 | 31.88 | 41.541 |
| B:Thr69 | -0.7 |  | Helix | 54.871 | 47.299 | 41.382 | 54.818 |
| B:Ser70 | -0.8 |  | Helix | 1.949 | 1.446 | 1.717 | 2.388 |
| B:Phe71 | 2.8 |  | Helix | 11.038 | 7.548 | 5.519 | 5.068 |
| B:Val72 | 4.2 |  | Helix | 52.906 | 31.7 | 37.622 | 36.052 |
| B:Lys73 | -3.9 | 10.4 | Helix | 118.615 | 78.568 | 60.995 | 55.517 |
| B:Leu74 | 3.8 |  | Helix | 14.584 | 5.535 | 9.614 | 5.57 |
| B:Leu75 | 3.8 |  | Coil | 7.132 | 0 | 4.702 | 0 |
| B:Pro76 | -1.6 |  | Coil | 71.076 | 34.216 | 55.874 | 36.51 |
| B:Leu77 | 3.8 |  | Coil | 42.635 | 31.7 | 28.105 | 31.899 |
| B:Arg81 | -4.5 | 12 | Sheet | 53.188 | 52.685 | 23.187 | 29.86 |
| B:Tyr82 | -1.3 | 10 | Sheet | 4.025 | 4.025 | 1.833 | 2.389 |
| B:Ala83 | 1.8 |  | Sheet | 3.019 | 3.019 | 3.015 | 6.557 |
| B:Leu84 | 3.8 |  | Sheet | 7.013 | 6.541 | 4.623 | 6.582 |
| B:Tyr85 | -1.3 | 10 | Sheet | 27.475 | 27.475 | 12.514 | 16.305 |
| B:Asp86 | -3.5 | 3.9 | Sheet | 16.086 | 15.615 | 11.028 | 16.492 |
| B:Ala87 | 1.8 |  | Sheet | 2.516 | 1.006 | 2.513 | 2.186 |
| B:Asp98 | -3.5 | 3.9 | Sheet | 41.725 | 38.234 | 28.605 | 40.383 |
| B:Leu99 | 3.8 |  | Sheet | 27.164 | 19.121 | 17.907 | 19.241 |
| B:Val100 | 4.2 |  | Sheet | 1.51 | 1.51 | 1.073 | 1.717 |
| B:Phe101 | 2.8 |  | Sheet | 4.025 | 4.025 | 2.013 | 2.703 |
| B:Ile102 | 4.5 |  | Sheet | 10.064 | 10.064 | 6.298 | 8.869 |
| B:Phe103 | 2.8 |  | Sheet | 19.121 | 19.121 | 9.56 | 12.838 |
| B:Trp104 | -0.9 |  | Sheet | 10.064 | 10.064 | 4.156 | 5.26 |
| B:Ala105 | 1.8 |  | Coil | 11.254 | 6.541 | 11.24 | 14.208 |
| B:Pro110 | -1.6 |  | Coil | 73.919 | 63.904 | 58.109 | 68.188 |
| B:Leu111 | 3.8 |  | Helix | 149.994 | 132.839 | 98.879 | 133.671 |
| B:Ser113 | -0.8 |  | Helix | 47.371 | 36.573 | 41.745 | 60.421 |
| B:Lys114 | -3.9 | 10.4 | Helix | 56.285 | 53.393 | 28.943 | 37.728 |
| B:Met115 | 1.9 |  | Turn | 87.748 | 77.485 | 48.223 | 60.058 |
| B:Ile116 | 4.5 |  | Helix | 50.318 | 47.802 | 31.492 | 42.129 |
| B:Tyr117 | -1.3 | 10 | Helix | 28.577 | 26.693 | 13.015 | 15.841 |
| B:Ala118 | 1.8 |  | Helix | 27.355 | 22.643 | 27.323 | 49.18 |
| B:Lys126 | -3.9 | 10.4 | Helix | 114.362 | 74.945 | 58.808 | 52.957 |
| B:Lys127 | -3.9 | 10.4 | Helix | 110.264 | 72.373 | 56.701 | 51.14 |
| B:Phe128 | 2.8 |  | Coil | 29.088 | 21.133 | 14.544 | 14.189 |
| B:Thr129 | -0.7 |  | Coil | 128 | 99.614 | 96.534 | 115.45 |
| B:Gly130 | -0.4 |  | Coil | 45.94 | 24.153 | 64.136 | 100.524 |
| B:Gly155 | -0.4 |  | Coil | 20.782 | 13.083 | 29.013 | 54.45 |
| B:Asn156 | -3.5 |  | Coil | 130.111 | 110.53 | 88.149 | 114.338 |
| B:Val157 | 4.2 |  | Coil | 81.459 | 72.961 | 57.926 | 82.976 |
| B:Val158 | 4.2 |  | Coil | 40.734 | 17.611 | 28.966 | 20.029 |
| B:Val159 | 4.2 |  | Coil | 105.508 | 69.439 | 75.028 | 78.97 |
| B:Ser160 | -0.8 |  | Sheet | 28.697 | 23.162 | 25.289 | 38.266 |
| B:Leu161 | 3.8 |  | Sheet | 12.938 | 8.554 | 8.529 | 8.608 |
| B:Pro165 | -1.6 |  | Coil | 92.721 | 71.955 | 72.889 | 76.779 |
| B:Leu166 | 3.8 |  | Coil | 69.064 | 13.083 | 45.528 | 13.165 |

D. **Complex 9**

| **Residue name** | **Hydrophobicity** | **pKa** | **Secondary structure** | **Residue solvent accessibility** | **Side chain solvent accessibility** | **% residue solvent accessibility** | **% side chain solvent accessibility** |
| --- | --- | --- | --- | --- | --- | --- | --- |
| A:Met1 | 1.9 |  | Coil | 143.22 | 42.837 | 78.708 | 33.203 |
| A:Ala2 | 1.8 |  | Coil | 35.668 | 15.599 | 35.626 | 33.88 |
| A:Val7 | 4.2 |  | Sheet | 24.105 | 7.548 | 17.141 | 8.584 |
| A:Ile12 | 4.5 |  | Helix | 38.114 | 36.229 | 23.854 | 31.929 |
| A:Lys13 | -3.9 | 10.4 | Turn | 95.637 | 84.2 | 49.179 | 59.497 |
| A:Val14 | 4.2 |  | Helix | 33.553 | 28.681 | 23.86 | 32.618 |
| A:Phe15 | 2.8 |  | Helix | 7.548 | 7.548 | 3.774 | 5.068 |
| A:Ile29 | 4.5 |  | Helix | 76.124 | 60.885 | 47.643 | 53.659 |
| A:Ala35 | 1.8 |  | Sheet | 3.019 | 3.019 | 3.015 | 6.557 |
| A:Val36 | 4.2 |  | Sheet | 5.032 | 5.032 | 3.578 | 5.722 |
| A:Leu37 | 3.8 |  | Sheet | 5.032 | 5.032 | 3.317 | 5.063 |
| A:Phe38 | 2.8 |  | Sheet | 0 | 0 | 0 | 0 |
| A:Cys39 | 2.5 | 9 | Sheet | 5.702 | 5.702 | 4.391 | 7.366 |
| A:Leu40 | 3.8 |  | Sheet | 17.06 | 1.006 | 11.246 | 1.013 |
| A:Ser41 | -0.8 |  | Coil | 22.196 | 20.686 | 19.56 | 34.175 |
| A:Gln46 | -3.5 |  | Sheet | 104.687 | 104.687 | 58.713 | 83.511 |
| A:Ile47 | 4.5 |  | Sheet | 3.77 | 0 | 2.359 | 0 |
| A:Ile48 | 4.5 |  | Sheet | 70.948 | 70.948 | 44.404 | 62.528 |
| A:Val49 | 4.2 |  | Sheet | 50.733 | 25.662 | 36.077 | 29.185 |
| A:Glu50 | -3.5 | 4.3 | Sheet | 44.768 | 44.768 | 25.383 | 36.272 |
| A:Gln54 | -3.5 |  | Sheet | 59.81 | 35.745 | 33.544 | 28.514 |
| A:Ile55 | 4.5 |  | Sheet | 16.605 | 15.095 | 10.392 | 13.304 |
| A:Leu56 | 3.8 |  | Coil | 103.646 | 93.591 | 68.325 | 94.177 |
| A:Val57 | 4.2 |  | Coil | 98.544 | 67.426 | 70.075 | 76.681 |
| A:Gly58 | -0.4 |  | Coil | 51.794 | 16.605 | 72.308 | 69.11 |
| A:Asp59 | -3.5 | 3.9 | Coil | 53.888 | 33.65 | 36.944 | 35.541 |
| A:Val64 | 4.2 |  | Coil | 41.764 | 36.229 | 29.699 | 41.202 |
| A:Tyr68 | -1.3 | 10 | Helix | 16.469 | 16.469 | 7.501 | 9.774 |
| A:Thr69 | -0.7 |  | Helix | 31.405 | 20.566 | 23.685 | 23.836 |
| A:Ser70 | -0.8 |  | Helix | 16.653 | 9.081 | 14.675 | 15.003 |
| A:Phe71 | 2.8 |  | Turn | 1.006 | 1.006 | 0.503 | 0.676 |
| A:Val72 | 4.2 |  | Helix | 41.948 | 37.235 | 29.829 | 42.346 |
| A:Lys73 | -3.9 | 10.4 | Helix | 94.598 | 76.556 | 48.645 | 54.095 |
| A:Leu74 | 3.8 |  | Helix | 8.898 | 6.541 | 5.865 | 6.582 |
| A:Leu75 | 3.8 |  | Coil | 14.393 | 3.019 | 9.488 | 3.038 |
| A:Pro76 | -1.6 |  | Coil | 38.696 | 29.688 | 30.42 | 31.678 |
| A:Leu77 | 3.8 |  | Turn | 103.381 | 94.598 | 68.15 | 95.19 |
| A:Arg81 | -4.5 | 12 | Sheet | 47.798 | 47.798 | 20.838 | 27.091 |
| A:Tyr82 | -1.3 | 10 | Sheet | 2.5 | 0 | 1.139 | 0 |
| A:Ala83 | 1.8 |  | Sheet | 3.961 | 2.516 | 3.957 | 5.464 |
| A:Leu84 | 3.8 |  | Sheet | 1.414 | 0 | 0.932 | 0 |
| A:Tyr85 | -1.3 | 10 | Sheet | 26.525 | 26.525 | 12.081 | 15.741 |
| A:Asp86 | -3.5 | 3.9 | Sheet | 30.255 | 26.893 | 20.742 | 28.404 |
| A:Ala87 | 1.8 |  | Coil | 15.974 | 9.057 | 15.955 | 19.672 |
| A:Asp98 | -3.5 | 3.9 | Coil | 24.2 | 19.736 | 16.591 | 20.845 |
| A:Leu99 | 3.8 |  | Sheet | 12.418 | 7.548 | 8.186 | 7.595 |
| A:Val100 | 4.2 |  | Sheet | 2.013 | 2.013 | 1.431 | 2.289 |
| A:Phe101 | 2.8 |  | Sheet | 5 | 4.529 | 2.5 | 3.041 |
| A:Ile102 | 4.5 |  | Sheet | 4.529 | 4.529 | 2.834 | 3.991 |
| A:Phe103 | 2.8 |  | Sheet | 9.56 | 9.56 | 4.78 | 6.419 |
| A:Trp104 | -0.9 |  | Sheet | 5.032 | 5.032 | 2.078 | 2.63 |
| A:Ala105 | 1.8 |  | Coil | 12.891 | 10.064 | 12.876 | 21.858 |
| A:Pro110 | -1.6 |  | Coil | 57.33 | 48.305 | 45.068 | 51.544 |
| A:Leu111 | 3.8 |  | Helix | 111.822 | 97.617 | 73.715 | 98.228 |
| A:Ser113 | -0.8 |  | Helix | 0 | 0 | 0 | 0 |
| A:Lys114 | -3.9 | 10.4 | Helix | 51.526 | 51.526 | 26.496 | 36.409 |
| A:Met115 | 1.9 |  | Turn | 100.197 | 98.217 | 55.065 | 76.127 |
| A:Ile116 | 4.5 |  | Turn | 1.51 | 1.51 | 0.945 | 1.33 |
| A:Tyr117 | -1.3 | 10 | Turn | 0 | 0 | 0 | 0 |
| A:Ala118 | 1.8 |  | Turn | 45.047 | 25.662 | 44.993 | 55.738 |
| A:Lys126 | -3.9 | 10.4 | Helix | 146.075 | 121.946 | 75.116 | 86.169 |
| A:Lys127 | -3.9 | 10.4 | Turn | 118.39 | 105.571 | 60.879 | 74.598 |
| A:Phe128 | 2.8 |  | Turn | 0.503 | 0.503 | 0.252 | 0.338 |
| A:Thr129 | -0.7 |  | Turn | 77.362 | 67.618 | 58.345 | 78.367 |
| A:Gly130 | -0.4 |  | Turn | 40.22 | 20.63 | 56.151 | 85.864 |
| A:Gly155 | -0.4 |  | Coil | 19.648 | 11.07 | 27.43 | 46.073 |
| A:Asn156 | -3.5 |  | Coil | 153.998 | 126.661 | 104.333 | 131.025 |
| A:Val157 | 4.2 |  | Coil | 82.425 | 73.967 | 58.614 | 84.12 |
| A:Val158 | 4.2 |  | Coil | 34.225 | 12.076 | 24.337 | 13.734 |
| A:Val159 | 4.2 |  | Coil | 94.654 | 51.827 | 67.309 | 58.941 |
| A:Ser160 | -0.8 |  | Coil | 65.765 | 56.101 | 57.955 | 92.683 |
| A:Leu161 | 3.8 |  | Coil | 42.267 | 42.267 | 27.863 | 42.532 |
| A:Pro165 | -1.6 |  | Coil | 53.697 | 36.229 | 42.212 | 38.658 |
| A:Leu166 | 3.8 |  | Coil | 145.068 | 61.891 | 95.631 | 62.278 |
| B:Met1 | 1.9 |  | Coil | 10.909 | 6.038 | 5.995 | 4.68 |
| B:Ala2 | 1.8 |  | Coil | 29.575 | 26.165 | 29.54 | 56.831 |
| B:Val7 | 4.2 |  | Coil | 19.353 | 0.503 | 13.762 | 0.572 |
| B:Ile12 | 4.5 |  | Helix | 62.833 | 61.388 | 39.325 | 54.102 |
| B:Lys13 | -3.9 | 10.4 | Helix | 109.971 | 98.422 | 56.55 | 69.546 |
| B:Val14 | 4.2 |  | Helix | 17.044 | 15.599 | 12.12 | 17.74 |
| B:Phe15 | 2.8 |  | Helix | 32.187 | 31.197 | 16.094 | 20.946 |
| B:Ile29 | 4.5 |  | Helix | 27.172 | 27.172 | 17.006 | 23.947 |
| B:Ala35 | 1.8 |  | Sheet | 0 | 0 | 0 | 0 |
| B:Val36 | 4.2 |  | Sheet | 0.503 | 0.503 | 0.358 | 0.572 |
| B:Leu37 | 3.8 |  | Sheet | 0.503 | 0.503 | 0.332 | 0.506 |
| B:Phe38 | 2.8 |  | Sheet | 4.465 | 3.522 | 2.232 | 2.365 |
| B:Cys39 | 2.5 | 9 | Sheet | 14.524 | 14.524 | 11.185 | 18.763 |
| B:Leu40 | 3.8 |  | Sheet | 36.684 | 20.127 | 24.183 | 20.253 |
| B:Ser41 | -0.8 |  | Coil | 24.049 | 16.445 | 21.193 | 27.169 |
| B:Gln46 | -3.5 |  | Coil | 74.1 | 72.655 | 41.558 | 57.958 |
| B:Ile47 | 4.5 |  | Sheet | 0 | 0 | 0 | 0 |
| B:Ile48 | 4.5 |  | Sheet | 45.789 | 45.789 | 28.658 | 40.355 |
| B:Val49 | 4.2 |  | Sheet | 24.193 | 2.013 | 17.204 | 2.289 |
| B:Glu50 | -3.5 | 4.3 | Sheet | 79.4 | 78.393 | 45.019 | 63.516 |
| B:Gln54 | -3.5 |  | Sheet | 9.991 | 2.922 | 5.603 | 2.331 |
| B:Ile55 | 4.5 |  | Sheet | 0.471 | 0 | 0.295 | 0 |
| B:Leu56 | 3.8 |  | Coil | 62.897 | 62.394 | 41.463 | 62.785 |
| B:Val57 | 4.2 |  | Turn | 73.975 | 49.311 | 52.604 | 56.08 |
| B:Gly58 | -0.4 |  | Turn | 43.4 | 25.159 | 60.59 | 104.712 |
| B:Asp59 | -3.5 | 3.9 | Helix | 38.856 | 26.206 | 26.638 | 27.678 |
| B:Val64 | 4.2 |  | Coil | 71.906 | 67.426 | 51.133 | 76.681 |
| B:Tyr68 | -1.3 | 10 | Turn | 13.522 | 13.019 | 6.159 | 7.726 |
| B:Thr69 | -0.7 |  | Helix | 54.256 | 38.705 | 40.918 | 44.858 |
| B:Ser70 | -0.8 |  | Helix | 5.216 | 4.712 | 4.596 | 7.785 |
| B:Phe71 | 2.8 |  | Helix | 5.032 | 5.032 | 2.516 | 3.378 |
| B:Val72 | 4.2 |  | Helix | 25.686 | 18.618 | 18.266 | 21.173 |
| B:Lys73 | -3.9 | 10.4 | Turn | 112.866 | 75.662 | 58.039 | 53.464 |
| B:Leu74 | 3.8 |  | Turn | 15.942 | 8.554 | 10.509 | 8.608 |
| B:Leu75 | 3.8 |  | Coil | 10.12 | 3.522 | 6.671 | 3.544 |
| B:Pro76 | -1.6 |  | Coil | 86.827 | 46.292 | 68.256 | 49.396 |
| B:Leu77 | 3.8 |  | Coil | 73.704 | 62.394 | 48.587 | 62.785 |
| B:Arg81 | -4.5 | 12 | Sheet | 38.656 | 38.656 | 16.852 | 21.909 |
| B:Tyr82 | -1.3 | 10 | Sheet | 1.006 | 0 | 0.458 | 0 |
| B:Ala83 | 1.8 |  | Sheet | 3.003 | 2.516 | 2.999 | 5.464 |
| B:Leu84 | 3.8 |  | Sheet | 1.51 | 1.51 | 0.995 | 1.519 |
| B:Tyr85 | -1.3 | 10 | Sheet | 5.535 | 5.535 | 2.521 | 3.285 |
| B:Asp86 | -3.5 | 3.9 | Sheet | 6.126 | 6.126 | 4.2 | 6.47 |
| B:Ala87 | 1.8 |  | Sheet | 1.006 | 1.006 | 1.005 | 2.186 |
| B:Asp98 | -3.5 | 3.9 | Sheet | 67.865 | 59.911 | 46.527 | 63.278 |
| B:Leu99 | 3.8 |  | Sheet | 27.178 | 17.108 | 17.916 | 17.215 |
| B:Val100 | 4.2 |  | Sheet | 7.044 | 3.019 | 5.009 | 3.433 |
| B:Phe101 | 2.8 |  | Sheet | 6.317 | 1.006 | 3.158 | 0.676 |
| B:Ile102 | 4.5 |  | Sheet | 5.503 | 5.032 | 3.444 | 4.435 |
| B:Phe103 | 2.8 |  | Sheet | 10.567 | 10.567 | 5.283 | 7.095 |
| B:Trp104 | -0.9 |  | Sheet | 6.541 | 6.541 | 2.701 | 3.419 |
| B:Ala105 | 1.8 |  | Coil | 12.26 | 7.548 | 12.245 | 16.393 |
| B:Pro110 | -1.6 |  | Coil | 64.91 | 57.866 | 51.027 | 61.745 |
| B:Leu111 | 3.8 |  | Helix | 129.905 | 117.241 | 85.636 | 117.975 |
| B:Ser113 | -0.8 |  | Helix | 29.487 | 17.1 | 25.985 | 28.251 |
| B:Lys114 | -3.9 | 10.4 | Helix | 54.437 | 52.552 | 27.993 | 37.134 |
| B:Met115 | 1.9 |  | Helix | 2.013 | 2.013 | 1.106 | 1.56 |
| B:Ile116 | 4.5 |  | Helix | 11.573 | 11.573 | 7.243 | 10.2 |
| B:Tyr117 | -1.3 | 10 | Helix | 28.258 | 28.258 | 12.87 | 16.77 |
| B:Ala118 | 1.8 |  | Helix | 28.194 | 13.586 | 28.161 | 29.508 |
| B:Lys126 | -3.9 | 10.4 | Turn | 143.493 | 93.374 | 73.788 | 65.979 |
| B:Lys127 | -3.9 | 10.4 | Turn | 92.366 | 63.9 | 47.497 | 45.152 |
| B:Phe128 | 2.8 |  | Coil | 12.044 | 11.07 | 6.022 | 7.432 |
| B:Thr129 | -0.7 |  | Turn | 127.369 | 107.29 | 96.058 | 124.345 |
| B:Gly130 | -0.4 |  | Turn | 62.774 | 29.184 | 87.637 | 121.466 |
| B:Gly155 | -0.4 |  | Coil | 10.846 | 7.548 | 15.142 | 31.414 |
| B:Asn156 | -3.5 |  | Turn | 129.178 | 110.339 | 87.517 | 114.141 |
| B:Val157 | 4.2 |  | Turn | 80.98 | 80.509 | 57.585 | 91.559 |
| B:Val158 | 4.2 |  | Sheet | 22.939 | 5.032 | 16.312 | 5.722 |
| B:Val159 | 4.2 |  | Sheet | 72.299 | 37.235 | 51.412 | 42.346 |
| B:Ser160 | -0.8 |  | Sheet | 21.261 | 16.246 | 18.736 | 26.839 |
| B:Leu161 | 3.8 |  | Sheet | 28.05 | 10.567 | 18.491 | 10.633 |
| B:Pro165 | -1.6 |  | Coil | 106.235 | 76.483 | 83.513 | 81.611 |
| B:Leu166 | 3.8 |  | Coil | 209.091 | 100.636 | 137.836 | 101.266 |

**Table S5:** The table includes the APRs and their respective scores in cofilin-1 as predicted by the Aggrescan3D 2.0 server.

| **Residue number** | **Residue name** | **Aggrescan3D 2.0 score** |
| --- | --- | --- |
| 1 | M | 1.0589 |
| 2 | A | 0.1956 |
| 5 | V | 0.4394 |
| 6 | A | 0.1385 |
| 7 | V | 0.3835 |
| 14 | V | 0.0918 |
| 19 | K | 0.0685 |
| 20 | V | 0.9779 |
| 29 | V | 1.4531 |
| 39 | C | 0.0791 |
| 48 | I | 1.7539 |
| 49 | L | 1.3898 |
| 56 | L | 0.2323 |
| 63 | T | 0.0326 |
| 64 | V | 1.7175 |
| 69 | A | 0.0313 |
| 103 | F | 0.357 |
| 105 | A | 0.0094 |
| 110 | P | 0.0445 |
| 111 | L | 1.2329 |
| 115 | M | 0.2703 |
| 117 | Y | 0.2463 |
| 140 | Y | 0.8809 |
| 147 | C | 0.0961 |
| 157 | A | 0.3102 |
| 158 | V | 1.7454 |
| 166 | L | 0.9936 |
