## Supplementary material for "Computational and biochemical analyses reveal that cofilin-2 self assembles into amyloid-like structures and promotes the aggregation of other proteinaceous species: Pathogenic relevance to myopathies": Figures S1-S2

**Figure S1**

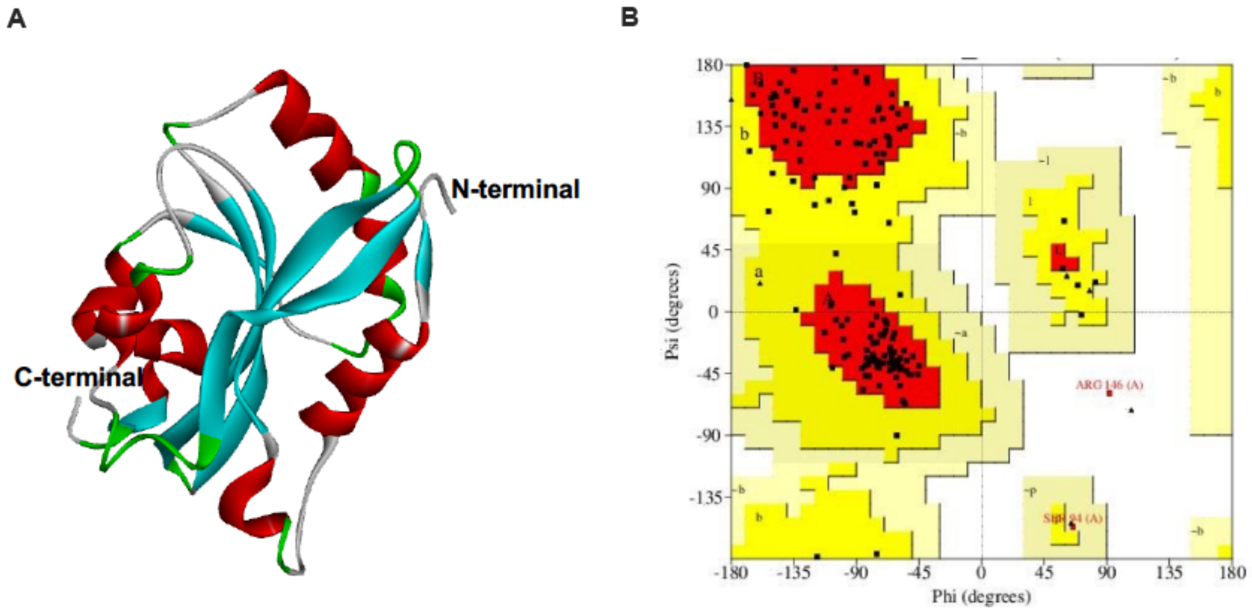


**Figure S1:** The cofilin-2 protein was modeled using the I-TASSER server, and the quality was then validated by PSVS (A) The cartoon representation of the best 3D model of cofilin-2 selected for the study. Cyan: β-sheets, red: α-helices, green: turns, and white: coils. (B) The Ramachandran plot of the modeled cofilin-2 structure.

Figure S2

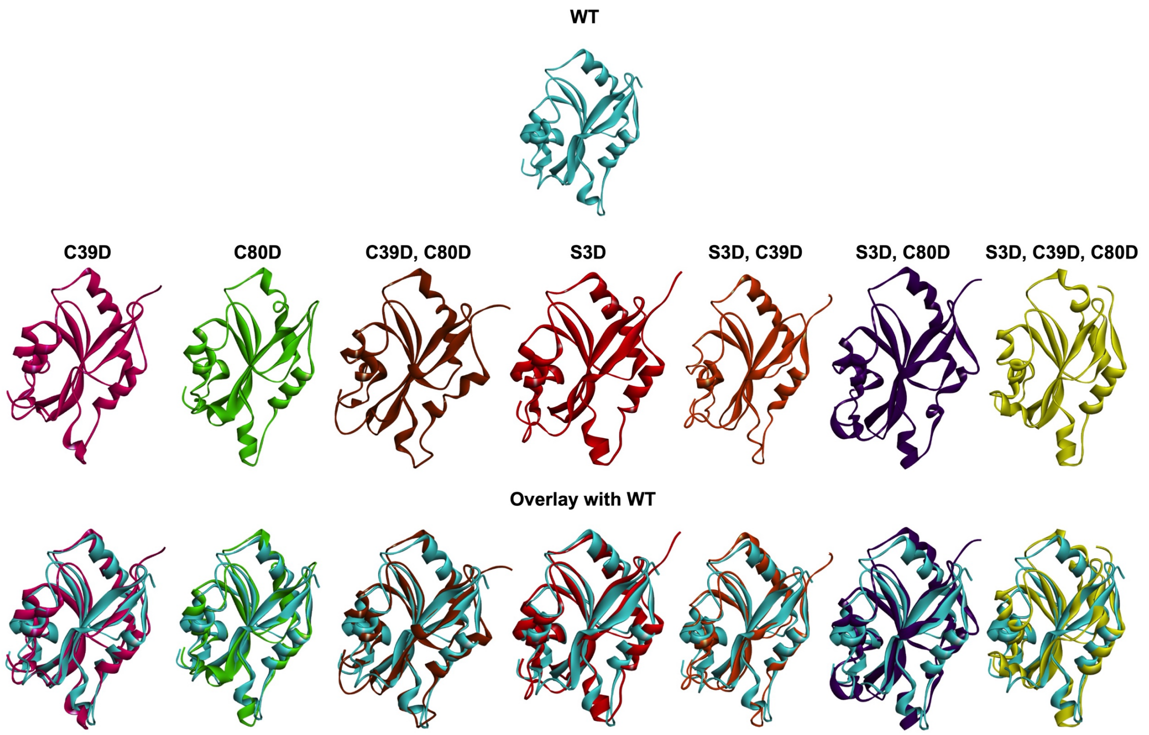


**Figure S2:** The 3D cartoon depictions of cofilin-2 (WT) and its mutants showing the alterations in the secondary structure caused by oxidation and phosphorylation mimics.
